# Microtubule plus-end regulation by centriolar cap proteins

**DOI:** 10.1101/2021.12.29.474442

**Authors:** Funso E. Ogunmolu, Shoeib Moradi, Vladimir A. Volkov, Chris van Hoorn, Jingchao Wu, Nemo Andrea, Shasha Hua, Kai Jiang, Ioannis Vakonakis, Mia Potočnjak, Franz Herzog, Benoît Gigant, Nikita Gudimchuk, Kelly E. Stecker, Marileen Dogterom, Michel O. Steinmetz, Anna Akhmanova

## Abstract

Centrioles are microtubule-based organelles required for the formation of centrosomes and cilia. Centriolar microtubules, unlike their cytosolic counterparts, grow very slowly and are very stable. The complex of centriolar proteins CP110 and CEP97 forms a cap that stabilizes the distal centriole end and prevents its over-elongation. Here, we used in vitro reconstitution assays to show that whereas CEP97 does not interact with microtubules directly, CP110 specifically binds microtubule plus ends, potently blocks their growth and induces microtubule pausing. Cryo-electron tomography indicated that CP110 binds to the luminal side of microtubule plus ends and reduces protofilament peeling. Furthermore, CP110 directly interacts with another centriole biogenesis factor, CPAP/SAS- 4, which tracks growing microtubule plus ends, slows down their growth and prevents catastrophes. CP110 and CPAP synergize in inhibiting plus-end growth, and this synergy depends on their direct binding. Together, our data reveal a molecular mechanism controlling centriolar microtubule plus-end dynamics and centriole biogenesis.

## Introduction

Centrioles are conserved organelles that play an important role in diverse processes such as cell division, motility, polarity and signaling. They are required for the assembly of centrosomes, the major microtubule (MT)-organizing centers in animal cells, and form the basal bodies of cilia and flagella (reviewed in ^1–5^). Defects in centriole components and centriole number have been linked to different human diseases, such as cancer, and to developmental disorders, including microcephaly and ciliopathies ^1–5^.

Centrioles are barrel-like structures, which typically contain nine MT triplets with the length in the range of several hundred nanometers. Centriole biogenesis relies on multiple specialized proteins, which set the nine-fold symmetry through a scaffolding structure, the cartwheel, and organize highly stable MT triplets and centriolar appendages ^3–5^. Unlike cytoplasmic MTs, which grow at a rate of 10-20 µm/min, centriolar MTs elongate with a rate of a few tens of nanometers per hour ^6–8^. This can be explained by the presence of specific centriolar factors that stabilize MTs and control their growth. Previous work has shown that the MT-binding centrosomal-P4.1-associated-protein (CPAP, or SAS-4 in worms and flies), which is essential for the formation of centriolar MTs and centriole elongation (reviewed in ^4, 9^), plays a role in preventing outgrowth of MT extensions from the distal centriole end ^10^. CPAP performs this function by capping MT plus ends through a specialized domain that binds to and occludes the surface of the tip-exposed β-tubulin ^10–12^. In vitro reconstitution experiments showed that CPAP tracks growing MT ends and stabilizes MTs by preventing catastrophes and making MT growth slow and persistent ^10^. However, these effects of CPAP on MT polymerization are not sufficient to explain how the elongation of centriolar MTs is restricted.

Another strong candidate for regulating centriolar MT plus-end growth is the “cap” structure observed at the distal ends of centrioles. The major components of this cap are CP110 and CEP97, which, similar to CPAP, regulate centriole elongation and prevent uncontrolled extension of the plus ends of centriolar MTs ^13–16^. The effects of CP110 and CEP97 on centriole length are species- and cell-specific. In mammalian cells, CP110 and CEP97 counteract the ability of CPAP to promote centriole elongation ^13, 14^. In different types of *Drosophila* cells and tissues, dependent on the cellular context, both elongation and shrinkage of centrioles were reported upon the loss of CP110 and CEP97 ^8, 17–20^. The emerging picture from these studies is that CP110 and CEP97 can counteract changes in centriole length imposed by well-studied positive or negative regulators of centriolar MT growth, such as CLASP or kinesin-13, respectively ^9, 19, 20^. CP110 and CEP97 are also required for early stages of cilia formation ^18, 21, 22^, but the cap structure that these proteins form needs to be removed from the basal body to allow the formation of axonemal MTs ^15, 23–25^.

While genetic and cell biological studies strongly support the role of CP110 and CEP97 in forming a regulatory cap at the distal centriolar end, biochemical understanding of their activities is limited. It is well established that the two proteins interact with each other and with a number of other factors involved in the biogenesis of centrioles and cilia ^9, 15, 26–31^. However, it is currently unknown whether and how CP110 and CEP97 interact with MTs and whether they exert autonomous or non-autonomous effects on MT growth. To fill in this knowledge gap, we reconstituted in vitro the activities of purified CP110 and CEP97 on dynamic MTs. We found that CP110 can specifically interact with MT plus ends and block their growth through its C-terminal domain, whereas CEP97 does not interact with MTs directly. We also found that CP110 can directly bind to CPAP, and that this interaction potentiates the plus-end-blocking activity of CP110. However, CP110 and CPAP do not interfere with each other’s activities if their binding interface is perturbed, suggesting that they associate with distinct sites on MT plus ends. Cryo-electron tomography data further indicated that CP110 interacts with the luminal side of MT plus ends and inhibits protofilament peeling. Together, our data indicate the CP110 is a MT growth inhibitor whose activity can be modulated by other centriole and cilia assembly factors.

## Results

### CP110 binds to MT plus ends and blocks their growth

In order to investigate the direct effects of CP110 and CEP97 on MT growth, we used in vitro reconstitution assays, in which MTs polymerizing from GMPCPP-stabilized seeds attached to a glass slide are observed by Total Internal Reflection Fluorescence (TIRF) microscopy ^10, 32^. Full-length CP110 and CEP97 with an N-terminal GFP tag were purified from HEK293 cells (Figure 1a, Supp. Figure S1a,b). We observed that GFP-CP110 could bind to the plus ends of seeds and block their elongation at concentrations above 30 nM, whereas MT minus ends, which grow more slowly than the plus ends, were not affected (Figure 1b,c, Supp. Figure S1c). At concentrations lower than 30 nM, GFP-CP110 could occasionally bind to MT plus ends and induce pausing followed by catastrophes (Figure 1c). In contrast, CEP97-GFP displayed no binding to MTs and no effect on their dynamics at concentrations up to 50 nM (Supp. Figure S1d). The addition of up to 240 nM CEP97 to the assays with 30 nM CP110 had no effect on MT seed blocking (Supp. Figure S1e). Unfortunately, in all these assays, we observed significant aggregation of CP110, which complicated quantitative analyses, and the addition of CEP97 did not solve this problem. We have also tried to generate CP110 deletion mutants, but they were even more difficult to purify, and their biochemical activities therefore could not be tested.

**Figure 1.**
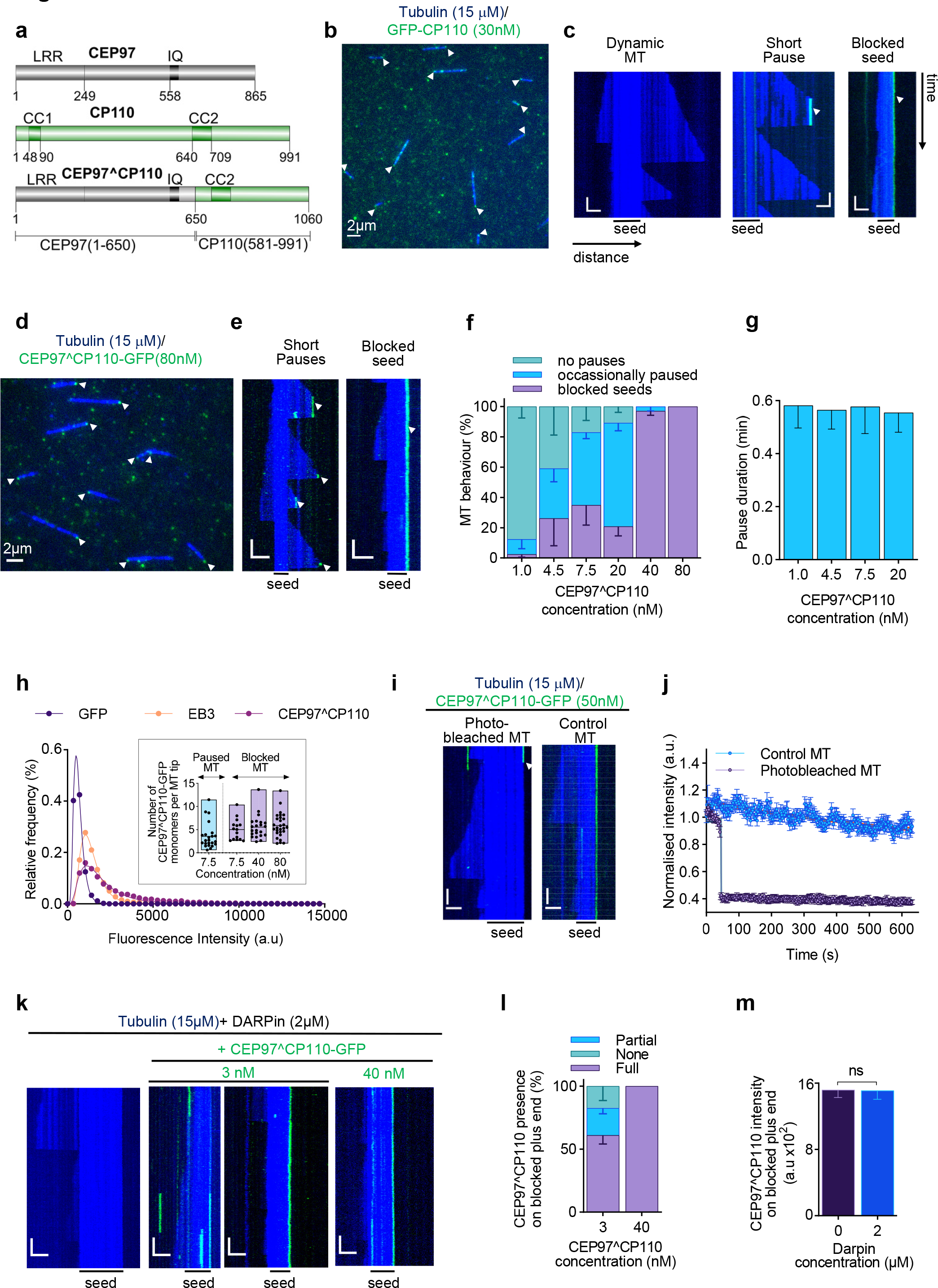
CP110 binds to MT plus ends and blocks their growth. **(a)** A scheme of the domain organization of human CEP97, CP110 and the CEP97^CP110 fusion protein. **(b,d)** Still images from time-lapse movies of GFP-CP110 (b) or CEP97^CP110-GFP (d) (green) blocking growth of GMPCPP-stabilized MT seeds (blue) in vitro. Arrows point to blocked MT plus ends. **(c,e)** Representative kymographs showing MT growth with 15 μM tubulin alone (“dynamic MT”), or in the presence of GFP-CP110 (c) or CEP97^CP110 (e), causing short pauses or blocking the plus ends of GMPCPP seeds. Bars, 2 µm (horizontal); 1 min (vertical). (**f)** The proportion of MTs with no pauses observed over a 10 min period, occasionally paused or fully blocked with increasing CEP97^CP110-GFP concentrations. The number of analyzed MTs was 156, 122, 133, 106, 184, 193 at 1.0, 4.5, 7.5, 20, 40 and 80 nM CEP97^CP110-GFP, respectively; n = 3 independent assays except for 1.0 nM where n=5. **(g)** The mean MT pause duration at CEP97^CP110-GFP concentrations inducing short pauses. n = 18, 45, 102, 133 pausing events at 1.0, 4.5, 7.5, and 20 nM CEP97^CP110-GFP, respectively from 3 independent assays. **(h)** Histograms of fluorescence intensities at the initial moment of observation of single molecules of the indicated proteins immobilized on coverslips (symbols) and the corresponding fits with lognormal distributions (lines); 6865, 14082 and 6942 molecules for GFP (monomers), EB3 (dimers) and CEP97^CP110, respectively. The inset shows the plot of the number of CEP97^CP110-GFP molecules causing short pausing or blocking of MT growth. The values were obtained by comparing the fitted mean intensity of CEP97^CP110-GFP at MT tips with the fitted mean intensity of single GFP molecules in parallel chambers. Floating bars represent maximum to minimum intensities of CEP97^CP110-GFP molecules relative to GFP per condition, with the line showing the mean value. A total of 15, 22, 28 fully blocked MTs were analyzed at CEP97^CP110-GFP concentration of 7.5, 40 and 80 nM, while 23 paused MTs were analyzed at 7.5 nM. **(i)** Recovery dynamics of CEP97^CP110-GFP at MT plus ends after photobleaching. The moment of photobleaching is indicated by a white arrowhead. Bars are the same as in panel (c). **(j)** Quantification (mean ± SEM) of CEP97^CP110-GFP signal in photobleaching experiments. n = 28 photobleached MTs, n = 12 for control MTs from 3 independent experiments. **(k)** Representative kymographs of MT plus end growth in the presence of DARPin (TM- 3)2 alone or together with CEP97^CP110. Bars, 2 µm (horizontal); 1 min (vertical). **(l, m)** Presence of CEP97^CP110-GFP (3 nM or 40 nM) at MT plus ends (l) and mean fluorescence intensities of CEP97^CP110-GFP (40 nM) at blocked MT plus end (m) in the presence or absence of DARPin (TM-3)2. (l) The number of analyzed MTs was 91 for 3nM CEP97^CP110-GFP and 110 for 40 nM CEP97^CP110-GFP, n = 2 and 4 independent experiments, respectively. (m) n = 76 and 83 MTs at 0 µM and 2 µM DARPin (TM-3)2, respectively; error bars are mean ± SEM; ns – no significant differences in mean at P<0.05, (Mann–Whitney test).

It is known that the N-terminus of CP110 binds to the middle part of CEP97 ^15^. To improve the protein quality, we tested the idea that a CEP97-CP110 chimera, in which some of the domains of both proteins were omitted and the remaining parts fused together, could lead to a well-behaved protein. We initially screened different chimeric proteins by their localization in U2OS cells. We found that a protein containing residues 1-650 of CEP97 and residues 581-991 of CP110 (termed here CEP97^CP110, Figure 1a) displayed a clear centriole localization and used it for subsequent experiments. As shown in Figure 1d-f, GFP-tagged CEP97^CP110 could potently block MT seed elongation in vitro, similar to full-length CP110, but was less aggregation-prone (Figure 1d-f). While this protein did not bind along MT shafts, it could specifically bind to MT plus ends and completely block their growth at concentrations exceeding 40 nM, while MT minus ends, which could be distinguished by their slower polymerization rate, underwent normal dynamics. At lower concentrations of CEP97^CP110-GFP, MTs could still grow from both ends, but the binding of the chimera caused transient plus end pausing with an average duration of ∼0.6 min, followed by MT depolymerization (Figure 1d-g). These results demonstrate that our CP110-CEP97 fusion approach provides a way to study the effect of CP110 on MT dynamics.

Next, we used measurements of fluorescence intensity to determine how many molecules of the CEP97^CP110 chimera are sufficient to block MT growth. By comparing the intensity of individual GFP-tagged CEP97^CP110 molecules immobilized on glass to the intensity of single molecules of purified GFP (monomers) or GFP-EB3 (dimers), we found that CEP97^CP110-GFP is a dimer (Figure 1h). We then compared the intensity of CEP97^CP110-GFP blocking or pausing a MT tip to the intensity of individual molecules of the same protein immobilized on glass in a separate chamber. We found that, on average, four CEP97^CP110-GFP molecules (two dimers) were observed at MT ends undergoing transient pausing at 7.5 nM, and six CEP97^CP110-GFP molecules (three dimers) were seen at the fully blocked tips of the seeds at concentrations between 7.5 and 80 nM (Figure 1h). The total number of CEP97^CP110-GFP molecules bound to the MT plus end rarely exceeded 10 monomers, which is lower than the number of protofilaments present in GMPCPP-stabilized MTs that predominantly contain 14 protofilaments ^33^. To determine the dynamics of CEP97^CP110-GFP on blocked MT plus ends, we used Fluorescence Recovery after Photobleaching (FRAP) and found that the protein displays no turnover (Figure 1i,j). Taken together, these data suggest that a relatively small number of CEP97^CP110-GFP molecules (fewer than the number of MT protofilaments) is sufficient to arrest MT plus-end growth, and that they do so by stably binding to MT tips. Since CEP97 does not associate with MTs on its own, this binding depends on the C-terminal half (residues 581-991) of CP110.

The most obvious way for a protein to block MT plus-end growth is by occluding the longitudinal interface of β-tubulin and prevent α-tubulin from binding to it. An agent known to have such an activity is the tubulin-specific designed ankyrin repeat protein (DARPin), which binds to β- tubulin and inhibits subunit addition to the plus end ^34, 35^. To gain additional insight into the mode of action of CP110, we tested the potential competition between CEP97^CP110-GFP and DARPin by using (TM-3)2, a dimeric version of the high affinity DARPin TM-3 ^35, 36^. At 2 µM, the (TM-3)2 DARPin completely blocked elongation of MT plus ends, but not minus ends, in the presence of 15 µM soluble tubulin (Figure 1k). However, CEP97^CP110-GFP could still efficiently bind to such blocked MT plus ends even when present at a 3 nM concentration (Figure 1k, l). We also found no difference in the intensity of 40 nM CEP97^CP110-GFP at the MT plus ends in the presence or absence of 2 µM DARPin (Figure 1m). These data indicate that CEP97^CP110 binds to non-dynamic MT plus ends in the presence of a very large molar excess of two types of molecules, tubulin dimers and DARPin, which have a strong affinity for the plus-end-exposed part of β-tubulin.

### CP110 binds to MT plus ends from the luminal side and reduces protofilament peeling

To get further insight into the binding of the CEP97^CP110 chimera to MT plus ends and its effect on MT tip structure, we turned to cryo-electron tomography (cryoET). We reconstructed 3D volumes containing MTs grown in the presence or absence of 80 nM CEP97^CP110-GFP. Samples were frozen after 5-20 min of incubation of GMPCPP seeds with 15 µM soluble tubulin at 37°C. While we could not determine whether individual MT ends were growing or shortening, we assumed that the majority of MTs must be elongating, because in our *in vitro* assays, the time MTs spend growing is much longer than the time they spend shortening. The use of a recently developed denoising algorithm (^37^, see Methods for data processing details), allowed us to enhance the signal-to-noise ratio in the reconstructed 3D volumes, and to significantly improve the segmentation of individual protofilaments at the ends of MTs and their manual tracing. As reported previously ^38^, most MT ends in our samples terminated with curved protofilaments (Figure 2a).

**Figure 2.**
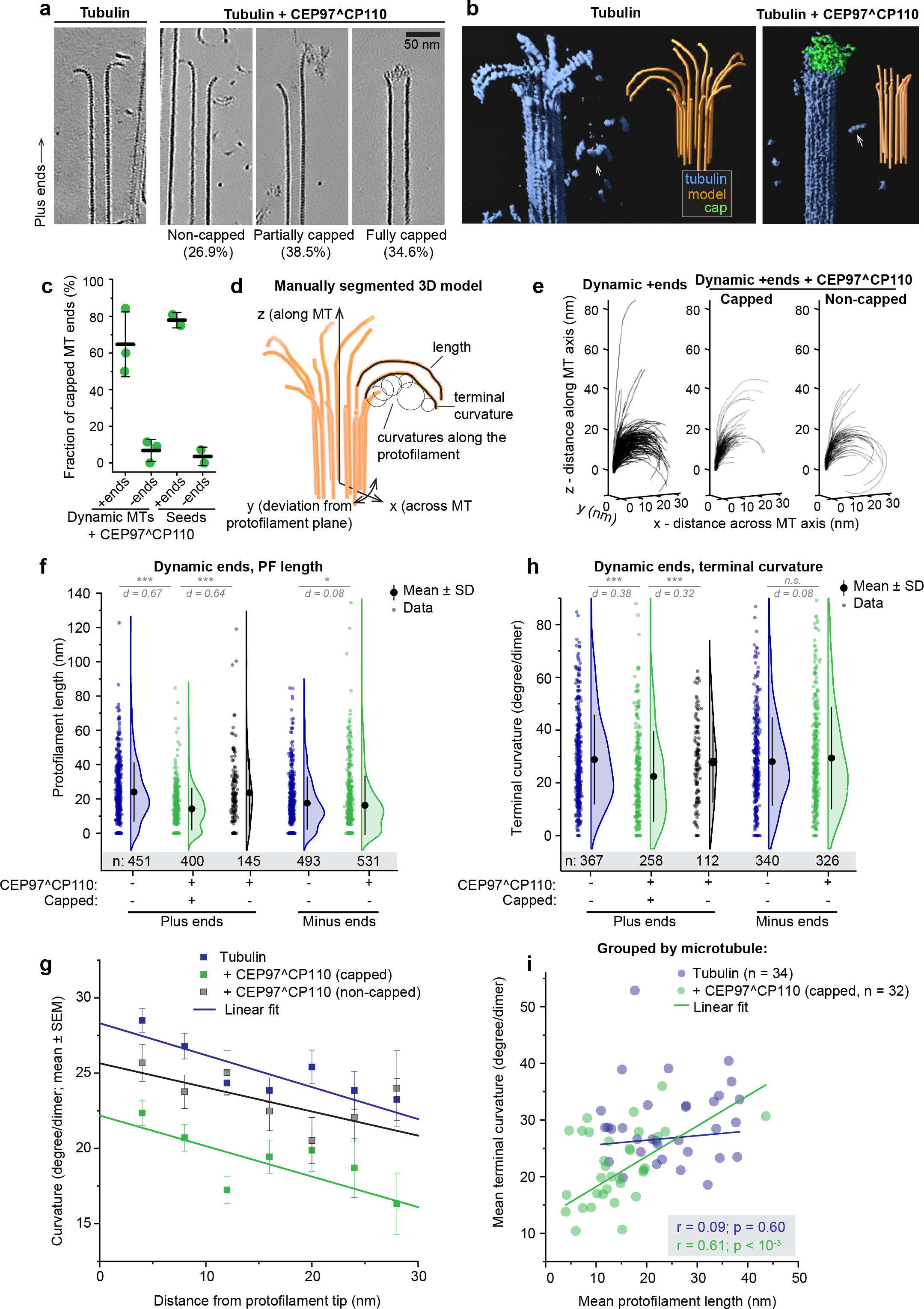
CEP97^CP110 forms caps at MT plus ends and straightens their protofilaments. (**a**) Slices through denoised tomograms containing MT plus ends in the absence or presence of 80 nM CEP97^CP110. (**b**) Segmented and 3D rendered volumes containing MT plus ends (blue), capping density (green) and manually segmented 3D models tracing protofilament shapes (orange). Arrows point to soluble tubulin oligomers. (**c**) Fraction of MT ends associated with a capping density. Data points: individual grids, line: mean ± SD. (**d**) Parameters extracted from manual segmentations of terminal protofilaments. (**e**) All protofilament traces obtained from plus ends in the presence of soluble tubulin, aligned at their origin. (**f**) Distribution of protofilament lengths for samples imaged in the presence of soluble tubulin. Here and below: shown are individual data points (dots), mean (circle) and SD (error bars). Statistical summary: d indicates effect size (Cohen’s d) expressed in units of SD; ***, P< 0.001; *, P<0.05 (Mann-Whitney test). (**g**) Average curvature of protofilaments aligned at their distal tips. Error bars show SEM, straight lines are the results of linear fitting. (**h**) Distribution of terminal curvature of protofilaments with non-zero length, obtained in the presence of soluble tubulin. Statistical summary: d indicates effect size (Cohen’s d) expressed in units of SD; ns – no significant difference; ***, P< 0.001 (Mann-Whitney test). (**i**) Correlation between average terminal curvature and average protofilament length per MT plus end. *r* – Pearson correlation coefficient, *p* – probability that the slope of the correlation is different from zero.

As expected, MT growth from GMPCPP-stabilized seeds produced primarily 14- protofilament MTs (170 out of 202; 84%), which allowed for unambiguous polarity determination of most MT ends (Supp. Figure S2a,b) ^39^. Interestingly, in the presence of CEP97^CP110, we observed ‘caps’ at MT ends, which were attached to a subset of protofilaments (partially capped) or blocking the whole MT lumen (fully capped) (Figure 2a, b, Supp. Figure S2b, Supp. Video S1). Capping densities were observed much more frequently at MT plus ends (Figure 2c): 78% of plus ends carried a cap (38 out of 52), compared to only 9% of capped minus ends (5 out of 56). Some MT plus ends were attached to larger structures, which we also considered as full caps (Supp. Figure S2b). Out of three sample preparations with soluble tubulin and CEP97^CP110, two were prepared with CEP97^CP110 added after the tubulin mix was subjected to high-speed centrifugation, and this led to the presence of large structures presumably formed by the chimeric protein (see Supp. Figure S2b for examples). In the sample with CEP97^CP110 added to the tubulin mix before centrifugation, we still observed caps predominantly at plus ends (50% capped plus ends, 9% capped minus ends); however, no full caps were seen in this sample. Therefore, fully capped MTs in our assays likely carry many more copies of CEP97^CP110 than determined by our TIRF assays (Figure 1h), which were performed after centrifugation of the tubulin-CEP97^CP110 mix. We also prepared samples with GMPCPP seeds and CEP97^CP110 without soluble tubulin and found that in these conditions, the assembly of large CEP97^CP110 structures was more prominent (see Figure S2b for examples). However, we observed a similar MT capping frequency: 78% of GMPCPP seeds in the absence of soluble tubulin were capped or attached end-on to large structures (38 out of 49) compared to 3% of capped minus ends (1 out of 30) (Figure 2c). Importantly, most caps appeared to interact with the luminal side of the protofilaments (Figure 2a, b, Supp. Figure S2b, Supp. Video S1) To determine whether CEP97^CP110-mediated capping affected protofilament shapes at MT ends, we manually traced protofilaments in tomograms (Figure 2b,d). From these manually segmented 3D models we obtained protofilament length (measured from the first segment bending away from the MT cylinder) and curvature along the protofilament (Figure 2d). Contrary to a previous report ^38^, protofilaments in our samples frequently deviated from their planes (Supp. Figure S2c). This difference forced us to modify the previously reported analysis to account for the full 3D coordinates of terminal protofilaments (Figure 2e, see Methods for details).

The presence of a CEP97^CP110 cap correlated with shorter protofilaments at dynamic MT plus ends; protofilaments at non-capped MT ends in the presence of the protein were not different from those imaged in its absence (Figure 2f). Since statistical analysis frequently yielded significant but tiny differences between sets of hundreds of individual protofilaments, we used Cohen’s d as a measure of effect size, and only regarded differences characterized by d > 0.2 x Standard Deviation (SD) as biologically significant. Minus ends in the absence of CEP97^CP110 had slightly shorter protofilaments than plus ends, but the presence of CEP97^CP110 had only a very minor effect on their length (Figure 2f, d < 0.2). Similarly, average protofilament curvature was reduced at CEP97^CP110-capped plus ends, but not at uncapped plus ends or minus ends in the presence of CEP97^CP110 (Supp. Figure S3a). As reported previously, protofilaments became more curved as they deviated from the MT wall (Figure 2g) ^38^. The presence of CEP97^CP110 reduced the average curvature of the terminal protofilament segments for capped plus ends, but not for uncapped plus or minus ends (Figure 2h).

Since CEP97^CP110 blocked MT growth at the seed in our TIRF experiments (Figure 1), we wondered whether the changes we observed in the shapes of the protofilaments were CEP97^CP110-mediated, or simply reflected the difference between a growing MT end with a GTP cap and a stable GMPCPP-stabilized end of the seed. To address this question, we analyzed the structures of GMPCPP-stabilized seed ends with and without CEP97^CP110. In the absence of soluble tubulin, GMPCPP-stabilized seeds still depolymerized, so we repeated this experiment in presence of a low concentration of tubulin to protect the seeds without elongating them (Supp. Figure S3b). In both conditions, with and without a protective concentration of tubulin, the presence of CEP97^CP110 caps correlated with straightened plus-end protofilaments (Supp. Figure 3c, Supplementary Table 1).

**Figure 3.**
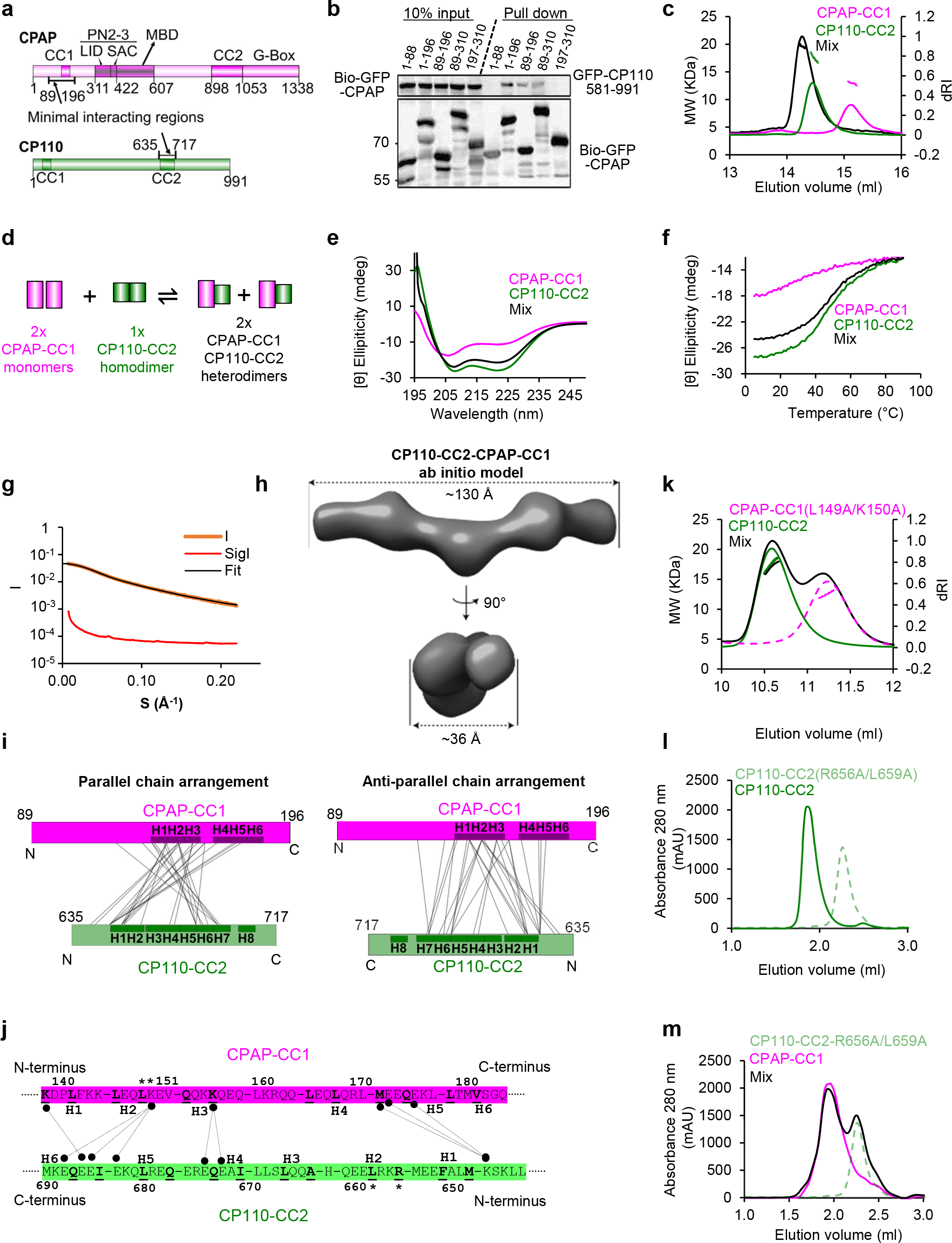
Structural and biophysical characterization of the CPAP-CP110 interaction. **(a)** Schematic representation of the domain organization of human CPAP and CP110. Numbers indicate amino acid positions. The minimal regions CPAP and CP110 that interact with each other are indicated. The domain nomenclature is as follows: CC, coiled coil; MBD, MT-binding domain; PN2-3, tubulin-binding PN2-3 domain; G-box, glycine-rich C-terminal domain. **(b)** Streptavidin pull- down assays with BioGFP-CPAP truncations as bait and GFP-CP110 (581-99) as prey. The assays were performed with the extracts of HEK293T cells co-expressing the constructs and BirA and analyzed by Western blotting with anti-GFP antibodies. **(c)** SEC-MALS analysis of CPAP-CC1 (magenta lines), CP110-CC2 (green lines) and an equimolar mixture of CPAP-CC1 and CP110-CC2 (black lines). **(d)** Proposed reaction mechanism for CPAP-CC1 and CP110-CC2 association. **(e, f)** CD spectra (e) recorded at 15°C and thermal unfolding profiles (f) recorded by CD at 222 nm. Proteins and colors as in (c). **(g, h)** SAXS analysis of the CPAP-CC1/CP110-CC2 heterodimer. **(g)** Solution X-ray scattering intensity over scattering angle from a 1:1 mixture (monomer equivalents) of CPAP-CC1 and CP110-CC2. The fit to the data yielding the interatomic distance distribution is shown with a black line. **(h)** Surface representation of the X-ray scattering volume of CPAP-CC1- CP110-CC2, at 32 ± 3 Å estimated precision, derived from averaging 22 particle models calculated by *ab initio* fit to the scattering data. **(i, j)** Chemical crosslinking followed by mass spectrometry analysis of CPAP-CC1-CP110-CC2. (**i**) Schematic representations of parallel (left) and antiparallel (right) arrangements of CPAP-CC1 and CP110-CC2 chains in the CPAP-CC1/CP110-CC2 heterodimer. Predicted heptad repeats (H) in each chain are indicated. Observed chemical crosslinks between residues of CPAP-CC1 and CP110-CC2 are indicated by thin lines. **(j)** Normalized inter- chemical crosslinks observed between CPAP-CC1 and CP110-CC2 in the CPAP-CC1/CP110-CC2 heterodimer. The heptad **<u>a</u>** and **<u>d</u>** position residues are shown in bold and are underlined. The CPAP- CC1 and CP110-CC2 residues that were mutated in this study are highlighted with asterisks. **(k)** SEC- MALS analysis of CPAP-CC1 L149A/K150A (magenta dashed lines), CP110-CC2 (green solid lines) and an equimolar mixture of CPAP-CC1 L149A/K150A and CP110-CC2 (black solid lines). **(l, m)** Analytical SEC analysis of CPAP-CC1 and CP110-CC2 variants. **(l)** Analytical SEC analysis of CP110-CC2 (green solid lines) and CP110-CC2 R656A/L659A (light green dashed lines). **(m)** Analytical SEC analysis of CPAP-CC1 (magenta lines), CP110-CC2 R656A/L659A (light green dashed lines), and an equimolar mixture of CPAP-CC1/CP110-CC2 R656A/L659A (black solid line).

Finally, we examined the correlation between protofilament length and curvature. Terminal curvature at free MT plus ends did not correlate with protofilament length. However, in the presence of CEP97^CP110-mediated caps, MT plus ends carrying shorter protofilaments were also characterized by reduced protofilament curvature (Figure 2i). Such a positive correlation was mainly observed at partially capped MT ends, because fully capped ends showed no correlation between average length and curvature of protofilaments (Supp. Figure S3d). We hypothesize that partially capped MT ends present a heterogeneous group that contains intermediate states between long, curved protofilaments, as observed at free MT plus ends, and short straight protofilaments as observed at MT plus ends fully capped by CEP97^CP110. We conclude that CEP97^CP110 reduces peeling of the terminal protofilaments at MT plus end, to which it likely binds from the luminal side.

### CP110 directly binds to CPAP

Having established that CP110 binds to the outermost MT plus end, we next wondered about its interplay with CPAP, a centriolar biogenesis factor which can also directly associate with protofilament termini at MT plus ends ^40^. The potential interaction between the two proteins has been suggested by proximity mapping ^26^, and here we tested whether the binding is direct. We co-expressed in HEK293T cells full-length CP110 and CPAP or their fragments tagged with either GFP alone or GFP and a biotinylation (Bio) tag together with biotin ligase BirA and performed streptavidin pull-down assays ^41^. We found that human full-length CP110 indeed associated with human full-length CPAP (Supp. Fig S4a-e). The C-terminal region 581-991 of CP110, which contains a predicted coiled-coil domain (CP110-CC2), was sufficient for the interaction with the full-length CPAP (Supp. Figure S4a, b). A shorter C-terminal CP110 fragment 581-700 still bound to CPAP, albeit weaker than longer fragments (Sup. Figure S4a,d). Further, we found that the N-terminal part of CPAP mediates the binding to full-length CP110 (Sup. Fig S4d,e) and that the CPAP fragment 89-196 including its predicted coiled-coil domain (CPAP-CC1) is sufficient for the association with CP110 581-991 (Figure 3a,b).

Next, we sought to analyze the interaction between N-terminal CPAP and C-terminal CP110 fragments in more detail using biophysical and structural methods. The recombinant expression and purification of CPAP 89-196 was straightforward, and we also produced a fragment containing residues 635-717 of CP110, which included the CP110-CC2 domain. From here onwards, the two fragments are referred to as CPAP-CC1 and CP110-CC2. The oligomerization state of these two domains as well as their combination was tested using size-exclusion chromatography coupled with multi-angle light scattering (SEC-MALS). For CPAP-CC1, these experiments revealed a single elution peak, corresponding to a molecular mass of 13.0 ± 1.8 kDa, consistent with the presence of a monomer (calculated mass of the monomer: 12.5 kDa). In contrast, CP110-CC2 revealed a single elution peak corresponding to a molecular mass of 17.5 ± 1.0 kDa, consistent with the formation of a homodimer (calculated mass of the monomer: 10.0 kDa). When the two proteins were mixed together in equimolar ratio, a single peak corresponding to a molecular mass of 19.7 ± 1.1 kDa was found, suggesting the formation of a CPAP-CC1/CP110-CC2 heterodimer (Figure 3c). Increasing the CPAP-CC1 concentration in the mixture by 2- and 3-fold supported this conclusion (Supp. Figure S5a,b). These results suggest that two CPAP-CC1 monomers react with one CP110-CC2 dimer to form two stable heterodimers in solution (Figure 3d).

### CP110 and CPAP interact by forming an anti-parallel coiled coil

Next, we analyzed the structure of CPAP-CC1, CP110-CC2, and CPAP-CC1/CP110-CC2 by circular dichroism (CD) spectroscopy. The far-UV CD spectrum of CPAP-CC1 recorded at 15°C, with minima at 220 and 205 nm, was characteristic of proteins displaying a mixture of helical and random-coil secondary structure content. In contrast, CP110-CC2 and a 1:1 mixture of CPAP-CC1 and CP110-CC2 (monomer equivalents) revealed CD spectra characteristic of mostly α-helical proteins, with minima at 208 and 222 nm (Figure 3e). The stability of the proteins was subsequently tested by thermal unfolding profiles monitored by CD at 222 nm. CPAP-CC1 revealed a broad, non-cooperative unfolding profile characteristic of a largely unfolded protein, whereas CP110-CC2 and a 1:1 mixture (monomer equivalents) of CPAP-CC1 and CP110-CC2 revealed sigmoidal and cooperative unfolding profiles characteristic of well-folded, α-helical coiled-coil proteins (Figure 3f). These results suggest that CPAP-CC1 is largely unfolded while CP110-CC2 and a mixture of CPAP- CC1 and CP110-CC2 forms α-helical coiled-coil structures.

To assess whether the CPAP-CC1/CP110-CC2 complex forms a canonical, extended coiled coil and to further probe the dimerization of CP110-CC2, we performed SEC coupled with small angle X-ray scattering (SEC-SAXS) experiments. Following buffer subtraction, the SAXS data were consistent with the presence of a monodisperse species in solution (Figure 3g, Supp. FigureS5c) with a radius of gyration, Rg, of 3.5 nm as estimated by Guinier approximation. To gain insight into the overall shape of CPAP-CC1/CP110-CC2 and CP110-CC2 in solution, we derived the pairwise distance distribution function, P(r), of these molecules (Figure 3h, Supp. Figure S5d), which suggested the presence of elongated particles in both cases, with a maximum dimension (interatomic distance, Dmax) of approximately 12.5 nm. This value is consistent with the calculated length of ∼12.0 nm for a two-stranded α-helical coiled coil of ∼80 amino acids. Accordingly, ab initio SAXS models derived from the P(r) distribution were consistent with the formation of extended coiled coils by CPAP-CC1/CP110-CC2 and CP110-CC2 (Figure 3h, Supp. Figure S5d).

To assess the orientation of the two chains in the CPAP-CC1/CP110-CC2 coiled-coil heterodimer, we performed chemical crosslinking coupled with mass spectrometry. To this end, the zero-length cross-linker DMTMM was used, a reagent that couples primary amines (side chain of lysines) with carboxylic acids (side chains of aspartate and glutamate) ^42^. We found 38 inter-crosslinks between CPAP-CC1 and CP110-CC2. By normalizing the intensities of the inter-links to the intra-links and ranking them accordingly ^43^, we selected the nine most abundant inter-links (Figure 3j), which together with our CD results suggested that CPAP-CC1 and CP110-CC2 form an antiparallel coiled-coil structure when mixed together (Figure 3i,j).

### Design of mutations that disrupt CPAP-CC1/CP110-CC2 coiled-coil formation

To test the functional relevance of the CPAP-CP110 interaction, we sought to create mutants that fail to associate. To this end, we mutated several conserved residues occupying either the predicted heptad **a** and **d** core positions and/or the **e** and **g** flanking positions of the predicted coiled-coil regions (Figure 3j). We found that simultaneous mutation of L149 and K150 at the heptad positions **d** and **e** of the second heptad repeat of CPAP-CC1 to alanines (CPAP-CC1 L149A/K150A) disrupted CPAP-CC1/CP110-CC2 heterodimer formation; notably, K150 was among the final ranking of crosslinked residues identified in our crosslinking experiments (Figure 3j). SEC-MALS experiments of CPAP- CC1 L149A/K150A yielded an elution peak corresponding to a molecular mass of 12.5 ± 0.5 kDa, similar to wild type CPAP-CC1 (Figure 3k, Supp. Figure S5e). Analysis of a 1:1 mixture of CPAP- CC1 L149A/K150A and CP110-CC2 (monomer equivalents) revealed two elution peaks, which corresponded to molecular masses of 16.7 ± 0.4 kDa (CP110-CC2 homodimer) and 13.0 ± 0.5 kDa (CPAP-CC1 L149A/K150A monomer), respectively (Figure 3k).

We further found that mutating R656 and L659 at the heptad position **a** and **d** of the second heptad repeat of CP110-CC2 to alanines (CP110-CC2-R656A/L659A) disrupts both CP110-CC2 homodimer as well as CPAP-CC1/CP110-CC2 heterodimer formation. Analytical SEC (aSEC) of CP110-CC2 R656A/L659A yielded a single elution peak, which corresponded to the elution of a monomeric protein (Figure 3l). Consistent with this finding, CD experiments with CP110-CC2- R656A/L659A revealed a spectrum with minima at around 220 and 205 nm and a broad, non-cooperative unfolding profile (Supp. Figure S5f, g). A subsequent aSEC analysis of a 1:1 mixture of CPAP-CC1 and CP110-CC2 R656A/L659A (monomer equivalents) revealed two elution peaks corresponding to monomers of CPAP-CC1 and CP110-CC2-R656A/L659A, respectively (Figure 3m).

Taken together, these results demonstrate that residues at key positions of the heptad repeats of both CPAP-CC1 and CP110-CC2 coiled-coil domains are critical for mediating CP110-CC2 homo- and CPAP-CC1/CP110-CC2 heterodimer formation. In combination with the results from pull-down experiments and the fact that coiled-coil domains behave as autonomous folding units ^44^, our results suggest that CPAP and CP110 can directly bind to each other and that their interaction can be disrupted by mutations in their respective coiled-coil domains.

### CPAP potentiates MT-blocking activity of CP110

Having devised a way to perturb the interaction between CP110 and CPAP, we set out to test its functional significance by using in vitro experiments. Since our previous work has shown that full-length CPAP does not behave well in vitro ^10^, we have generated a fusion of the N-terminal 1-607 fragment of CPAP to a dimer-forming leucine zipper of GCN4 and mCherry (Figure 4a, Supp. Figure S6). CPAPWT encompasses the two tubulin/MT-binding domains of CPAP, PN2-3 and MBD, which were part of the CPAPmini used in our previous study ^10^, but also contains the complete N-terminal region including the CP110-binding CC1 domain, which was absent in CPAPmini. In addition to wild type CPAP, we have also generated a similar fusion bearing the L149A/K150A mutations (CPAPmut) (Figure 4a, Supp. Figure S6). Similar to CPAPmini, both the wild type and mutant CPAP versions tracked growing MT plus ends (Figure 4b), displayed similar accumulation at the MT tips (Figure 4c), and imparted slow and processive MT plus-end growth with parameters that were similar to those previously described for CPAPmini ^10^ (Figure 4d). These data indicate that the CPAP-CC1 domain by itself does not contribute much to the MT plus-end regulation of CPAP.

**Figure 4.**
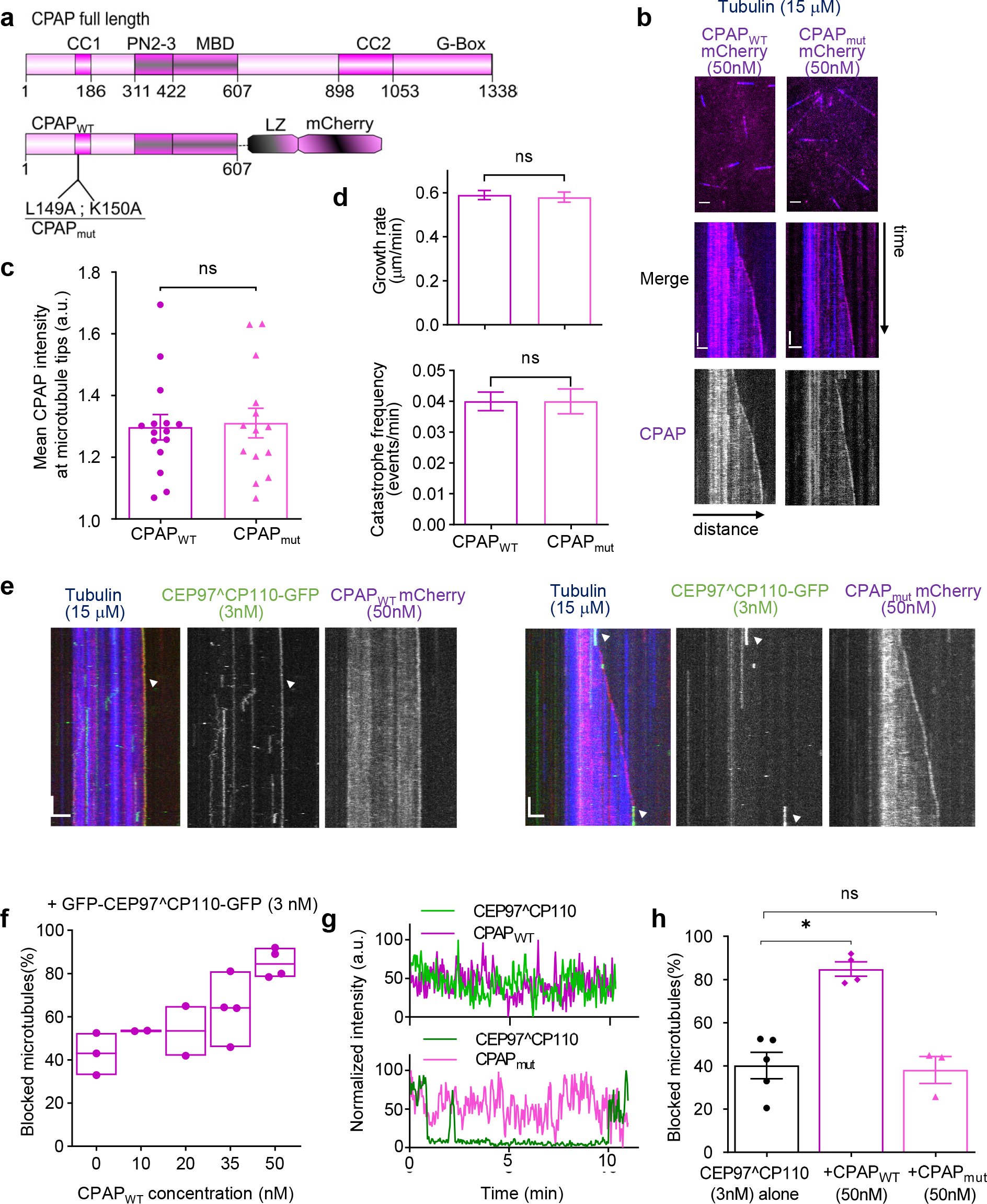
CPAP promotes the plus-end blocking activity of CP110. **(a)** A scheme of the domain organization of full-length human CPAP and truncated CPAP proteins, CPAPWT and CPAPmut with a leucine zipper (LZ) dimerization domain and an mCherry fluorescent tag at the C-termini. **(b)** Still images and representative kymographs of dynamic MTs growth in the presence of CPAP variants in vitro. Alexa 647 labelled MTs were grown from GMPCPP seeds in the presence of 50nM mCherry labelled CPAP variants - CPAPWT or CPAPmut. Bars, 2 µm (horizontal); 1 min (vertical). **(c)** Quantification of the CPAP intensities on growing MT plus tips in the presence of 50 nM of CPAP variants (n =15 MTs for CPAPWT and 14 MTs for CPAPmut from 2 independent experiments, respectively) **(d)** Growth rates and catastrophe frequencies of dynamic MTs in the presence of 50 nM CPAP variants (n= 151 and n=129 MT growth events for CPAPWT and CPAPmut from 4 and 5 independent experiments, respectively). For all plots, error bars are mean ± SEM; ns – no significant differences in mean at P<0.05, (Mann-Whitney test). **(e)** Representative kymographs illustrating MT dynamics in the presence of CEP97^CP110-GFP and CPAP variants. In overlays on the left, tubulin is shown in blue, CPAP in magenta and CEP97^CP110-GFP in green. White arrows point to blocked or paused plus end. Bars, 2 µm (horizontal); 1 min (vertical). **(f)** The effect of increasing CPAPWT concentration on MT plus end blocking by 3 nM CEP97^CP110-GFP. Each dot on the floating bars represents the percentage of blocked MTs from an independent experiment; lines are the median values from n independent experiments. The numbers of MTs are 192, 109, 116, 231 and 236 for the indicated CPAPWT concentrations. **(g)** The representative normalized intensity profile plot of CEP97^CP110-GFP (green) and CPAP variants (shades of magenta) on MTs plus ends during in vitro reconstitution with the respective proteins. **(h)** Each dot in the bar plots stands for an independent experiment; error bars are mean ± SEM, calculated based on n = 5, 4 and 3 experiments; 303, 236 and 153 MTs were analyzed, respectively; ns, no significant differences in mean; *, P<0.05 (Mann-Whitney test).

Next, we combined different concentrations of CPAPWT with 3 nM CEP97^CP110 (Figure 4e,f). At this low concentration, CEP97^CP110 by itself could block some but not all plus ends of the seeds; however, increasing concentrations of CPAPWT potentiated the blocking (Figure 4f), and the two proteins colocalized at MT plus ends (Figure 4g). In contrast, no increased seed blocking was observed when CPAPmut was used in these experiments (Figure 4h). Importantly, CPAPmut had no negative effect on the tip-blocking activity of CEP97^CP110, and the two proteins could still colocalize at MT plus ends (Figure 4g). These data suggest that at the concentrations tested, CPAP and CP110 do not compete with each other for the binding to MT tips and that their interaction makes MT growth inhibition more potent by stabilizing CP110 binding to MT plus ends or preventing tubulin addition to plus ends.

## Discussion

In this study, we have reconstituted in vitro the regulation of MT dynamics by the centriolar proteins CP110, CEP97, and CPAP. We showed that CP110 autonomously recognizes MT plus ends and inhibits their growth. These data are fully consistent with a large body of cell biological work showing that CP110 binds to distal ends of centrioles, prevents the overgrowth of centriolar MTs, and needs to be removed when centrioles are repurposed as ciliary basal bodies ^8, 13–20, 23–25, 45, 46^. In contrast, we found no evidence that CEP97 binds to MTs directly. This finding suggests that CEP97 affects centriolar MTs through other centriolar components, for example, through binding and regulation of CP110 ^9, 15, 46^. Our results obtained with CEP97^CP110 chimera are in line with this idea, as this fusion protein is better behaved in vitro than full-length CP110 or its fragments but is not more potent than CP110 alone. The MT-binding part of CP110 resides in its C-terminal part containing a dimeric coiled-coil domain that interacts with CPAP, as well as several putative helical and disordered regions, which, based on AlphaFold predictions ^47, 48^, are not expected to form folded protein domain(s). It is possible that the C-terminal part of CP110, besides its coiled-coil domain, assumes a stable structure only upon binding to MTs or other binding partners.

Our data provide important clues about the MT plus end-binding mechanism of CP110. First, CP110 stably binds to non-dynamic MT ends and can do so even in the presence of a 500-fold molar excess of soluble tubulin. This result suggests that the binding site of CP110 is specific for the MT lattice and may thus be formed by more than one tubulin subunit. Second, CP110 specifically blocks tubulin addition at MT plus ends, which suggests that it might occlude the longitudinal binding interface formed between β- and α-tubulin subunits from two different tubulin dimers. Notably, our data argue against a strong competition with two other proteins binding to the plus-end-exposed tip of β-tubulin, DARPin and CPAP, suggesting that the binding site of CP110 is distinct from those MT tip binders. Third, our cryoET data indicate that CP110 interacts with the luminal side of the MT plus end. One possibility is that CP110 interacts with the interface located between two adjacent protofilaments. Such a binding mode on the outside of the MT shaft is not unusual for proteins that specifically interact with MTs but not with soluble tubulin, such as End-Binding (EB) proteins ^49^, doublecortin ^50^, and CAMSAPs ^51^. CP110 might have some preference for the luminal interprotofilament groove at the plus end due to its potential asymmetry, because the protofilaments at MT ends can curl outwards, and at the plus ends, β-tubulins can separate somewhat further apart than α-tubulins. Such a mechanism would be analogous to the minus-end recognition by CAMSAP proteins ^51^. Alternatively, CP110 might bind along protofilaments, similar to MT inner proteins present in cilia ^52, 53^.

CP110 has two effects on MT plus-end dynamics: it can block the elongation of stable MT plus ends and induce their pausing, which means that it can suppress not only MT plus-end growth but also their shrinkage. Occlusion of the longitudinal interface of even a single β-tubulin subunit at the MT plus end has been shown to be sufficient to perturb MT growth ^54^. Catastrophe suppression of MT plus ends can also be caused by a small number of MT tip-bound molecules ^10, 55, 56^. It is thus not surprising that, similar to what we have previously observed in vitro for two other MT plus-end polymerization inhibitors, CPAP ^10^ and KIF21B ^55^, the number of CP110 molecules needed to strongly suppress MT elongation and induce pausing is much lower than the number of protofilaments: approximately 2-3 CP110 dimers were sufficient to block or pause MT growth from GMPCPP-stabilized MT seeds that contain on average 14 protofilaments ^33^. It thus appears that partial MT capping as observed in our cryoET data is sufficient to suppress both MT growth and depolymerization. In this context, the MT plus-end stabilizing effect of CP110 can be explained by its ability to suppress protofilament peeling, which could occur through either straightening the protofilaments or by enhancing their lateral interactions from the luminal side. It is also possible that the protofilament flaring, a feature that appears to be common for dynamic MT plus ends ^38, 57^, promotes tubulin addition and that blunt MT ends are more difficult to elongate. However, it is unlikely that changing protofilament shape alone is sufficient for the stable plus end blocking that we observe, and therefore, some steric occlusion of β-tubulin by CP110 at the outermost MT plus end seems likely. If steric occlusion occurs, it would inhibit the addition of tubulin dimers, which might result in shorter and less curved protofilaments at MT plus ends. Therefore, the difference in protofilament shapes induced by CP110 could be a consequence, rather than the cause of distinct tubulin on-rates.

We also found that CP110 directly interacts with CPAP and synergizes with it in inhibiting MT plus end elongation. CPAP contains several MT-binding domains, including the PN2-3 domain, which consists of LID and SAC subdomains that bind to the longitudinal interface and the outer surface of β-tubulin, respectively ^10–12^. The interaction between CP110 and CPAP is driven by the formation of a heterotypic antiparallel coiled-coil domain formed by CPAP-CC1 and CP110-CC2.

CPAP-CC1 is located N-terminally of the LID domain (Figure 3a), and the N-terminal part of the LID domain points towards the MT lumen ^12^. Therefore, CPAP-CC1 is expected to be ideally positioned to bind to CP110 at the luminal side of MT plus ends.

CP110 and CPAP synergize in plus-end blocking when they interact, but do not interfere with each other when their binding is perturbed. In our in vitro assays, CPAP tracks growing MT plus ends and recruits CP110; however, at the distal centriole ends, the CEP97-CP110 complex is likely maintained by additional interactions with other centriolar components and might recruit CPAP. It is possible that the synergy between CP110 and the LID domain of CPAP ensures efficient capping of centriolar MTs. In line with this view, the loss of the LID domain of CPAP does not abolish CPAP function in the formation of centriolar MTs but makes the centriolar cap structure permissive for MT overelongation ^10^. The availability of point mutations that specifically perturb the CPAP-CP110 interaction without interfering with MT binding opens the way to test the functional significance of their association in cells. Building complexity in the reconstitution system and increasing the resolution of cryoET analysis both in vitro and in intact centrioles will shed further light on the detailed mechanism of the centriolar cap attachment to MT ends.

## Methods

### Cell culture and transfection

HEK293T cells, obtained from the American Type Culture Collection (ATCC) Cells (HEK293T) were cultured in Dulbecco’s Modified Eagle’s Medium (DMEM), and Ham’s F10 (1:1) supplemented with 10% Fetal Calf Serum FCS and 5 U/ ml penicillin and 50 μg/ml streptomycin. Polyethylenimine (PEI, Polysciences) was used to transfect HEK293T cells for StrepTactin protein purification and streptavidin pull-down experiments. The cell line was routinely checked for mycoplasma contamination using the LT07-518 Mycoalert assay.

### Pull-down assays and Western blotting

For the pull-down assays, six-well plates with about 80-90% confluency were transfected with 1 µg plasmid DNA and 3 µL PEI (Polysciences) per well. Equal amount of the bait, prey, and BirA Biotin ligase DNA was used. One day after transfection, the medium was refreshed, and the second-day cells were harvested. Each sample was washed with ice-cold phosphate-buffered saline (PBS) and lysed on ice for 15 minutes with 100 µl lysis buffer (50 mM HEPES, 150 mM NaCl, 1% TritonX100) supplemented with protease and phosphatase inhibitor cocktail (Roche). 10% of the soluble fraction of the lysate was boiled with 4X sample buffer. Dynabeads® (Thermofisher) were blocked with 0.1% albumin from chicken egg white (Sigma) for 30 minutes and washed three times with wash buffer (50 mM HEPES, 150 mM NaCl, 0.1% Triton-X100, pH 7.4). The remaining 90% of the soluble fraction was incubated with the beads at 4°C for one hour while rolling. DynaMag-2 (Invitrogen) magnets were used for washing the beads. After three washes, the beads were boiled in a 2X sample buffer. All samples were loaded on SDS-PAGE gels, with a chosen percentage (6-9%) according to the protein size.

Unstained SDS gels were transferred to a nitrocellulose membrane using a semi-dry transfer cell (Bio-rad) for 2 hours at 12 volts. Membranes were blocked for 30 minutes with 2% BSA before adding the primary antibody to incubate overnight at 4°C. We used a rabbit polyclonal antibody against GFP (Abcam, ab290). The membrane was washed three times for five minutes in Phosphate-buffered saline containing 0.05% Tween-20 (PBST) before adding the secondary antibody. Goat antirabbit and goat anti-mouse InfraRedDye 800CW/680LT (Li-Cor Biosciences) secondary antibodies were used. After 1hour incubation, again three washes PBST were performed before imaging. Imaging was done on Odyssey CLx infrared imager (Li-Cor Biosciences).

### Protein expression and purification from HEK293T cells for in vitro reconstitution assays

Human CPAP and CP110 constructs were described previously ^10, 28^. To overexpress proteins in HEK293T cells, cDNAs of the human proteins were cloned into pTT5 based expression vectors (Addgene #52355). The constructs were tagged with Twin-Strep-tag (SII) and fluorescent proteins (GFP or mCherry):- SII-GFP-CP110, CEP97-GFP-SII, CEP97^CP110-GFP-SII (a chimera of N-terminal amino acids 1-650 from CEP97 and C-terminal 581-991 from CP110), CPAP607WT- mCherry-SII and CPAP607_mut-_mCherry-SII from CPAP full-length N-terminal amino acids 1-607 and purified from HEK293T cells using the StrepTactin affinity purification as previously described in ^10^. The cells were transfected with the plasmid DNA complexed at ratio 1:3 (w/w) with polyethyleneimine (1mg/mL) to form a PEI-DNA mixture in antibiotics-free Ham’s F-10 medium (Gibco) for 30 minutes at room temperature. The PEI-DNA mixture was afterwards gently added to the adherent HEK293T cells in complete DMEM and incubated at 37 °C in a 5% CO2 incubator. Cells were harvested two days post-transfection. The cells from one 15 cm dish were lysed in 500 μl lysis buffer (50 mM HEPES, 300 mM NaCl, 0.5% Triton X-100, pH 7.4) supplemented with protease inhibitors (Roche). After clearing debris by centrifugation, cell lysates were incubated with 10 μl StrepTactin beads (GE Healthcare) for 45 min. Beads were washed five times with lysis buffer without protease inhibitors and twice with wash buffer (50 mM HEPES, 150 mM NaCl, 0.05% Triton X-100). The proteins were eluted in 60 μl elution buffer (50 mM HEPES, 150 mM NaCl, 1mM MgCl2, 1 mM EGTA, 1 mM dithiothreitol (DTT), 2.5 mM d-Desthiobiotin and 0.05% Triton X-100, pH7.4). All purified proteins were snap-frozen in liquid nitrogen and stored at −80 °C.

### Mass spectrometry

To confirm the identity of purified proteins, purified protein samples were digested using S-TRAP microfilters (ProtiFi) according to the manufacturer’s protocol. Briefly, 4 µg of protein sample was denatured in 5% SDS buffer and reduced and alkylated using DTT (20 mM, 10 min, 95°C) and IAA (40 mM, 30 min). Next, samples were acidified, and proteins were precipitated using a methanol TEAB buffer before loading on the S-TRAP column. Trapped proteins were washed four times with the methanol TEAB buffer and then digested overnight at 37°C using 1ug Trypsin (Promega). Digested peptides were eluted and dried in a vacuum centrifuge before LC-MS analysis.

Samples were analyzed by reversed-phase nLC-MS/MS using an Ultimate 3000 UHPLC coupled to an Orbitrap Q Exactive HF-X mass spectrometer (Thermo Scientific). Digested peptides were separated using a 50 cm reversed-phase column packed in-house (Agilent Poroshell EC-C18, 2.7 µm, 50cm x 75 µm) and were eluted at a flow rate of 300 nl/min using a linear gradient with buffer A (0.1% FA) and buffer B (80% ACN, 0.1% FA) ranging from 13-44% B over 38 min, followed by a column wash and re-equilibration step. The total data acquisition time was 55 min. MS data were acquired using a DDA method with the following MS1 scan parameters: 60,000 resolution, AGC target equal to 3E6, maximum injection time of 20 msec, the scan range of 375-1600 m/z, acquired in profile mode. The MS2 method was set at 15,000 resolution, with an AGC target set to standard, an automatic maximum injection time, and an isolation window of 1.4 m/z. Scans were acquired using a fixed first mass of 120 m/z and a mass range of 200-2000, and an NCE of 28. Precursor ions were selected for fragmentation using a 1-second scan cycle, a dynamic exclusion time set to 10 sec, and a precursor charge selection filter for ions possessing +2 to +6 charges.

Raw files were processed using Proteome Discoverer (PD) (version 2.4, Thermo Scientific). MSMS fragment spectra were searched using Sequest HT against a human database (UniProt, year 2020) that was modified to contain protein sequences from our cloning constructs and a common contaminants database. The search parameters were set using a precursor mass tolerance of 20 ppm and a fragment mass tolerance of 0.06 Da. Trypsin digestion was selected with a maximum of 2 missed cleavages. Variable modifications were set as methionine oxidation, and protein N-term acetylation and fixed modifications were set to carbamidomethylation. Percolator was used to assign a 1% false discovery rate (FDR) for peptide spectral matches, and a 1% FDR was applied to peptide and protein assemblies. An additional filter requiring a minimum Sequest score of 2.0 was set for PSM inclusion. MS1 based quantification was performed using the Precursor Ion Quantifier node with default settings applied. Precursor ion feature matching was enabled using the Feature Mapper node. Proteins matching the common contaminate database were filtered out from the results table.

### Protein expression and purification from *E. coli* for biophysical and structural studies

CPAP-CC1 (residues 89-196) and CP110-CC2 (residues 635-717) were amplified by PCR and cloned into the bacterial expression vector PSPCm9 ^58^ containing N-terminal thioredoxin, a 6x His-tag and a PreScission cleavage site. Mutants of CPAP-CC1 and CP110-CC2 were generated using a PCR-based site-directed mutagenesis approach. The DNA sequences of all the established constructs were validated via sequencing.

Protein expression was performed in the *E. coli* strain BL21(DE3). In brief, LB medium containing 50 mg/ml of kanamycin was used for growing the transfected *E. coli* cells at 37°C. Once cell cultures reached an OD600 of 0.6, they were cooled down to 18°C and then induced with 0.4 mM isopropyl 1-thio-β-D-galactopyranoside (IPTG). Proteins were expressed overnight at 18°C. The next day, cells were harvested by centrifugation, washed in cold PBS buffer and lysed via sonication in a buffer containing 20 mM HEPES, pH 7.5, 1 M NaCl, 10% glycerol, 30 mM imidazole pH. 8.0, 5 mM β-mercaptoethanol, supplemented with protease cOmplete inhibitor cocktail (Roche) and DNAse (Sigma Aldrich). After high-speed centrifugation at 18,000g, the supernatants were collected and applied onto a HiTrap Ni-NTA column (Cytiva) for immobilized metal-affinity chromatography (IMAC) purification at 4°C. The bound proteins were washed extensively with IMAC buffer to remove non-specifically bound proteins. Bound proteins were eluted by increasing the concentration of imidazole to 500 mM. To cleave off the N-terminal thioredoxin-His fusion tag, the eluted fractions were pooled and incubated in the presence of His-tagged HRV 3C protease ^59^ overnight at 4°C in IMAC buffer. The cleaved samples were separated from non-cleaved proteins and HRV 3C protease via a HiTrap Ni-NTA purification step. Cleaved proteins were concentrated and loaded onto a size exclusion chromatography (SEC) HiLoad Superdex 75 16/60 column (Cytiva) for final purification in a buffer containing 20 mM HEPES, pH 7.5, 150 mM NaCl, 2 mM β-mercaptoethanol. The quality and identity of proteins were assessed by SDS-PAGE and mass spectrometry before storing at −80°C for further experiments.

### Circular dichroism (CD) spectroscopy

Far-UV CD spectra of proteins samples were recorded at 5°C using a Chirascan-Plus spectrophotometer (Applied Photophysics Ltd.), equipped with a computer-controlled Peltier element. A 400 µl of protein sample with the final concentration of 0.2 mg/ml in PBS was loaded into a quartz cuvette of 1 mm optical path length. The thermal stability of each protein sample was analyzed by monitoring their CD spectrum at 222 nm using constant heating from 5 to 85°C with 1°C min^-1^ intervals. The apparent midpoint of the transition, referring to the melting temperature, Tm, was determined by fitting the data points with the GraphPad Prism 7 by choosing the nonlinear least-square fitting function based on a sigmoid model.

### Size exclusion chromatography coupled with multi-angle light scattering (SEC-MALS)

SEC-MALS was done at 20°C using a Superdex S75 10/30 or a Superdex S200 10/30 column (Cytiva). The system was purged and equilibrated overnight using an Agilent UltiMate3000 HPLC in a buffer containing 20 mM HEPES, pH 7.5, 150 mM NaCl, 2 mM DTT with a flow rate of 0.5 ml/min. For each experiment, 15 µl of protein sample was loaded onto the respective SEC column at a concentration of ∼ 7 mg/ml. The molecular mass of protein samples was determined using the miniDAWN TREOS and Optilab T-rEX refractive index detectors (Wyatt Technology). For the data fitting, the Zimm model was selected in the ASTRA 6 software.

### SAXS data collection and analysis

SAXS data were collected at the small-angle scattering beamline B21 of the Diamond Light Source (Harwell, UK). Protein samples in 50 mM HEPES, pH 7.5, 100 mM NaCl, 1 mM DTT, 1 mM MgCl2 were passed through a Shodex (Munich, Germany) KW402.5-4F SEC column in line to the X-ray scattering measurement cell. Samples of 20 and 10 mg/ml protein concentrations and 80 μl volume were used; however, only data from the lower concentration samples were analyzed due to superior homogeneity as judged by the SEC profile. Buffer subtraction, summation of scattering intensities across peaks in size-exclusion chromatograms, calculation of the radius of gyration (Rg) from Guinier plots, estimation of molecular weight from scattering volume-of-correlation (Vc) plots, and evaluation of distance distribution functions (P(r)) were performed using Scatter3 ^60^. *Ab initio* calculation of molecular volumes from P(r) distributions was performed using DAMMIF ^61^. For each dataset, 23 bead-based models were derived using random starting seeds and assuming no internal volume symmetry (P1). Pairwise cross-correlation and averaging of models was performed by DAMAVER ^62^. The final CP110-CC2 envelope derives from averaging of 22 calculated models with NSD 0.67 ±0.05, while the CPAP-CC1/CP110-CC2 envelope is the average of 22 models with NSD 0.68 ± 0.04. Bead models were converted to volumetric envelopes using Situs 3 (ref ^63^); graphical representations were created in UCSF Chimera ^64^.

### Chemical crosslinking combined with mass spectrometry

CP110-CC2 homodimers and CPAP-CC1/CP110-CC2 heterodimers were crosslinked using an equimolar mixture of isotopically-labelled DSSG (H6/D6) (Di[Sulfosuccinimidyl]Glutarate, Creative molecules) for 20 min at 25°C and 1200 rpm at a final concentration of 0.5 and 1.25 mM, respectively. The reaction was quenched with ammonium bicarbonate at a final concentration of 100 mM for 10 min. Crosslinking was also performed using a zero-length crosslinker DMTMM (4-(4,6-dimethoxy- 1,3,5-triazin-2-yl)-4-methylmorpholinium chloride, Sigma Aldrich) for 6 min at 25 °C and 1200 rpm at a final concentration of 60 mM. The reaction was quenched using a desalting column (Thermo Scientific), followed by the addition of ammonium bicarbonate.

Crosslinked samples were denatured by adding 2 sample volumes of 8 M urea, reduced with 5 mM TCEP (Thermo Scientific) and alkylated by adding 10 mM iodoacetamide (Sigma-Aldrich) for 40 min at room temperature. Digestion was performed with lysyl endopeptidase (1:50 w/w, Wako) for 2 h followed by a second digest with trypsin at 35°C overnight at 1200 rpm (1:50 ratio w/w, Promega). Proteolysis was stopped by the addition of 1% (v/v) trifluoroacetic acid (TFA). Crosslinked peptides were purified by reversed-phase chromatography using C18 cartridges (Sep-Pak, Waters) and enriched on a Superdex Peptide PC 3.2/30 column (300 × 3.2 mm).

Fractions of crosslinked peptides were analyzed by liquid chromatography coupled to tandem mass spectrometry using an LTQ Orbitrap Elite (Thermo Scientific) instrument ^65^. Crosslinked peptides were identified using xQuest ^66^. The results were filtered with an MS1 tolerance window of −4 to 4 ppm and score ≥ 22 followed by manual validation. The intensities of the identified crosslinks were extracted and normalized by using a modified protocol of the previously published software xTract ^43^.

### In vitro reconstitution assay

The in vitro assays with dynamic MTs were performed under the same conditions as described previously by ^10^. Briefly, in vitro flow chambers for TIRF microscopy were assembled on microscopic slides by two strips of double-sided tape with plasma-cleaned glass coverslips. Flow chambers were functionalized by sequential incubation with 0.2 mg/ml PLL-PEG-biotin (Susos AG, Switzerland) and 1 mg/ml neutravidin (Invitrogen) in MRB80 buffer (80 mM piperazine-N, N[prime]-bis (2-ethane sulfonic acid), pH 6.8, supplemented with 4 mM MgCl2, and 1 mM EGTA). Afterwards, GMPCPP-stabilized MT seeds were attached to the coverslips through biotin– neutravidin interactions. The flow chambers were further blocked with 1 mg/ml κ-casein. The reaction mix containing the different concentrations and combinations of the respective purified proteins, MRB80 buffer supplemented with 14.5 µM porcine brain tubulin, 0.5 µM [X-rhodamine/Alexa 488/Alexa 647] labelled tubulin, 50 mM KCl, 1 mM GTP, 0.2 mg/ml κ-casein, 0.1% methylcellulose and oxygen scavenger mix [50 mM glucose, 400 µg ml^−1^ glucose oxidase, 200 µg/ml catalase and 4 mM DT], was added to the flow chamber after centrifugation in an Airfuge for 5 min at 119,000g. The flow chamber was sealed with vacuum grease, and dynamic MTs were imaged immediately at 30°C using a total internal reflection fluorescence (TIRF) microscope. All tubulin products were from Cytoskeleton Inc.

### Total internal reflection fluorescence (TIRF) microscopy

TIRF imaging was performed on a microscope set-up (inverted research microscope, Nikon Eclipse Ti-E), equipped with the perfect focus system (Nikon) and a Nikon CFI Apo TIRF 100/1.49 numerical aperture oil objective (Nikon). The microscope was supplemented with a TIRF-E motorized TIRF illuminator, modified by Roper Scientific/PICT-IBiSA Institut Curie, and a stage-top incubator (model no. INUBG2E-ZILCS, Tokai Hit) to regulate the temperature of the sample. Image acquisition was performed using either a Photometrics Evolve 512 EMCCD camera (Roper Scientific) or a Photometrics CoolSNAP HQ2 CCD camera (Roper Scientific) and controlled with MetaMorph7.7 software (Molecular Devices). The Evolve EMCCD camera’s final resolution was 0.066 µm/pixel, while with the CoolSNAP Myo CCD camera, it was 0.045 µm/pixel. For excitation lasers, we used 491 nm 100 mW Stradus (Vortran), 561 nm 100 mW Jive (Cobolt) and 642 nm 110 mW Stradus (Vortran). We used an ET-GFP 49002 filter set (Chroma) for imaging proteins tagged with GFP, an ET-mCherry 49008 filter set (Chroma) for imaging X-Rhodamine labelled tubulin or mCherry- tagged proteins, and an ET647 for imaging Alexa647 labelled tubulin. We used sequential acquisition for the imaging experiments.

### Analysis of MT plus end dynamics in vitro

Kymographs were generated using the ImageJ plugin KymoResliceWide v.0.4 (https://github.com/ekatrukha/KymoResliceWide). MT dynamics parameters were obtained from the kymographs. For the experiments determining the proportion of MTs blocked or paused, we manually observed the kymographs for the complete blocking, occasional pausing, and no visible effects on dynamic MTs by the added proteins. Values reported are fractions of the total MT population expressed in percentages. The quantitative data reported for each experiment were collected in at least two independent assays.

### Fluorescence recovery after photobleaching (FRAP) assay

The FRAP assay (bleaching of protein by a focused laser beam) of CEP97^CP110-GFP blocked MTs was done on the TIRF microscope equipped with an ILas system (Roper Scientific/PICT-IBiSA). In vitro MT, dynamics assay was performed in the presence of GMPCPP-stabilized MT seeds with 15 µM tubulin (supplemented with 3% rhodamine) and 80 nM CEP97^CP110-GFP. Photobleaching in the CEP97^CP110-GFP channel was performed with the 488-nm laser in regions with CEP97^CP110-GFP blocking MT plus end. In the case of control, no photobleaching was conducted.

### Single molecule counting and fluorescence intensity analysis

The single molecule counting and fluorescence intensity analysis was done as described in ^55^. Briefly, parallel flow chambers were made on the same plasma cleaned coverslip containing the appropriate dilutions of purified GFP, GFP-EB3 and CEP97^CP110-GFP in MRB80 buffer. After protein addition, the flow chambers were washed with MRB80 buffer, sealed with vacuum grease, and immediately imaged with a TIRF microscope. Images (about 40) of unexposed coverslip areas were acquired with 100 ms exposure time and low laser power. Single-molecule fluorescence spots were detected and fitted with 2D Gaussian function using custom-written ImageJ plugin DoM_Utrecht v.1.2.2 (https://github.com/ekatrukha/DoM_Utrecht). The fitted peak intensity values were used to build fluorescence intensity histograms. The histograms were fitted to Gaussian distributions using GraphPad Prism 9. To estimate the number of CEP97^CP110-GFP molecules that might be causing the observed pausing or blocking of MT plus end, we immobilized single molecules of GFP on the coverslip of one of the flow chambers. We performed the in vitro reconstitution assay with the different concentrations of CEP97^CP110-GFP in the adjacent chamber of the same coverslip. Images of single unbleached molecules were acquired first, while time-lapse imaging was performed on the in vitro assay using the same illumination parameters. The CEP97^CP110-GFP accumulations completely blocking or pausing dynamic MTs were manually located as regions of interest in each frame and fitted with 2D Gaussian as described above. For building the distributions of molecules at the MT tip, each CEP97^CP110-GFP intensity value at the MT plus end was normalized by the average GFP single molecules intensity from the adjacent chamber. We followed the same procedure described above in the instances where we compared the intensities of CPAP molecules and when we examined the influence of DARPin on CEP97^CP110 or CPAPWT at the plus end.

### CryoET sample preparation and microscopy

MTs were grown by incubating GMPCPP-stabilized, doubly cycled seeds, with 15 µM porcine brain tubulin (Cytoskeleton) in the polymerization buffer (80 mM K-PIPES pH 6.9, 4 mM MgCl2, 1 mM EGTA, 1 mM GTP, 1 mM DTT). The reaction mix was centrifuged in Beckman Airfuge for 5 min at 119,000g prior to mixing with seeds. In samples with CEP97^CP110-GFP present, 80 nM of the protein was added to the reaction mix before centrifugation (1 grid) or after centrifugation (2 grids). After incubation for 6-20 min at 37°C, 5 nm gold particles were added to the mix, and then 3.5 µl was transferred to a recently glow-discharged, lacey carbon grid suspended in the chamber of Leica EM GP2 plunge freezer, equilibrated at 37°C and 98% relative humidity. The grid was immediately blotted for 4 s and plunge-frozen in liquid ethane.

Images were recorded on a JEM3200FSC microscope (JEOL) equipped with a K2 Summit direct electron detector (Gatan) and an in-column energy filter operated in zero-loss imaging mode with a 30 eV slit. Images were recorded at 300 kV with a nominal magnification of 10000, resulting in a pixel size of 3.668 Å at the specimen level. Imaging was performed using SerialEM software ^67^, recording bidirectional tilt series starting from 0° ±60°; tilt increment 2°; total dose of 80-100 e−/Å2; target defocus -4 µm.

### 3D volume reconstruction and analysis

Tomographic tilt-series were processed as outlined in Suppl. Figure S7 (analysis flowchart). Direct electron detector movie frames were aligned using MotionCor2 ^68^ and then split into full, even and odd stacks. Tilt series alignment and tomographic reconstructions were performed on sums of full stacks with the IMOD software package using gold beads as fiducial markers ^69^. Final tomographic volumes were binned by two, corrected for contrast transfer function, and the densities of gold beads were erased in IMOD. 3D volumes were subsequently denoised using the cryoCARE procedure ^37^. For this, 3D reconstruction was performed on odd and even aligned stacks with the IMOD parameters identified for full stacks. We trained 2-3 denoiser models for each acquisition series and then applied one model to the rest of the tomograms in this series. Splitting of movie frames, reconstructing even and odd volumes, training data generation, model training and denoising was performed on a cluster of graphics processing units (GPU) using python scripts (available at https://github.com/NemoAndrea/cryoCARE-hpc04).

Subvolumes containing MT ends were manually extracted from denoised tomographic volumes and processed further. First, the polarity of MTs was determined on summed projections using moiré patterns of images Fourier-filtered at the origin using Fiji ^39, 70^. Following the previously published procedure to obtain protofilament coordinates ^38^, 3D models were manually built for each MT end in 3dmod ^69^. Each protofilament was stored as a separate contour, the first point in a contour was placed on a MT wall, the second point at the last segment of the protofilament that was still in the MT cylinder, and the following points were placed every 2-4 nm along the bending part of the protofilament. Accuracy of manual segmentation was constantly monitored in the Isosurface view of 3dmod, which contained both the rendered 3D representation of the tomographic volume and the manually built 3D model. This procedure resulted in 3D models such as those presented in Figure 2b,d. Coordinates of the protofilaments were then extracted using the ’howflared’ program in IMOD.

Protofilament coordinates were further analyzed using Matlab scripts available at https://github.com/ngudimchuk/Process-PFs. These scripts are based on the previously published ones ^38, 57^, but they were modified to account for protofilament shapes that deviated from 2D planes. As reported previously, the sampling along the protofilament was made uniform by interpolation and then smoothed using quadratic LOESS with a window of 10 points. Curvature was calculated as the angle between consecutive pairs of line segments in LOESS-smoothed traces.

To segment the denoised densities into ’tubulin’ and ’cap’, we used the *tomoseg* module of EMAN2.2 ^71^. Using a full denoised tomogram containing MTs grown in the presence of CEP97^CP110, we boxed reference regions sets containing (1) MT walls, (2) bent protofilaments at MT ends and soluble tubulin oligomers, (3) caps at MT ends and (4) ’bad’ regions containing carbon support, gold particles, ice contamination etc. These boxed sets were then manually segmented, and three neural networks were trained: 1 vs 4, 2 vs 4 and 3 vs 4. The resulting neural networks were applied to subvolumes containing MT ends, and the resulting segmentations were used to mask tomographic densities in UCSF Chimera ^64^. Masked densities were imported into Blender to make visualizations.

### Statistical analysis

Comparisons of protofilament shapes extracted from manually segmented tomograms: reported p-values are calculated using a Mann-Whitney test in OriginPro 9.0. Effect size is calculated as Cohen’s 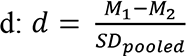d:, where *M1* and *M2* are means and pooled standard deviation 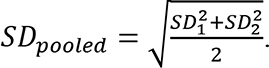 The reported p-values for the figures 1g, 1m, 4c, 4d and 4h were calculated using a two-tailed Mann-Whitney test in GraphPad Prism 9.

### Data and code availability

The data that support the conclusions are available in the manuscript; the original fluorescence microscopy datasets are available upon request to A.A. Tomography data presented in Figure 2ab and in Supplementary Video 1 are available from EMDB using the following accession codes: MTs in presence of tubulin and GMPCPP-stabilized seeds (EMD-14101 and EMD-14102), MTs in presence of CEP97^CP110, tubulin and GMPCPP-stabilized seeds (EMD-14103, EMD-14104 and EMD-14105). SAXS data and models are deposited in SASBDB: CPAP-CC1/CP110-CC2 heterodimer, accession code SASDNA3; CP110-CC2 homodimer, accession code SASDNB3.

Scripts used for data analysis are available at https://github.com/ekatrukha/KymoResliceWide, https://github.com/ekatrukha/DoM_Utrecht, https://github.com/NemoAndrea/cryoCARE-hpc04, and https://github.com/ngudimchuk/Process-PFs.

## Author Contributions

F.O. designed and performed protein purifications and in vitro reconstitution experiments, analyzed data and wrote the paper; S.M. designed and performed protein purification and biochemical and biophysical studies of protein-protein interactions, analyzed data and wrote the paper, V.A.V. designed and performed cryoET experiments, analyzed data and wrote the paper, C.H., J.W., S.H. and K.J generated protein fusions and performed biochemical experiments, I.V performed SAXS data collection and analysis, M.P. collected and analyzed the data on crosslinking followed by mass spectrometry and F.H. supervised these experiments, B. G. provided the (TM-3)2 protein, N.A and N.G generated algorithms and helped with processing of cryoET data, K.E.S. performed and analyzed mass spectrometry experiments, M.D., M.O.S and A.A. coordinated the project and wrote the paper.

## Acknowledgements

We thank R. Stucchi and A.F.M. Altelaar (Utrecht University) for the help with mass spectrometry experiments and W. Evers and A. Jakobi lab (TU Delft) for help with cryoET and valuable discussions. We also thank J. R. McIntosh (University of Colorado) for feedback on protofilament shape analysis and D. Mastronarde (University of Colorado) for modifying the ‘howflared’ program to extract 3D coordinates from protofilament models. This work was supported by the European Research Council Synergy grant 609822 to M.D. and A.A. and the Netherlands Organization for Scientific Research (NWO) ALWOP.440 grant to A.A, the Medical Research Council UK (MR/N009274/1) grant to I.V., EMBO Long Term Fellowship ALTF-840-2018 to F. O., Marie Curie COFUND Fellowship (Marie Skłodowska-Curie grant agreement 701647) to S. M., and grants from the Swiss National Science Foundation (31003A_166608 and 310030_192566 to M.O.S.). Analysis of the shapes of tubulin protofilaments was supported by the Russian Science Foundation grant 21- 74-20035 to N.G.

## Legends to Supplementary Figures and Video

**Supplementary Figure S1.**
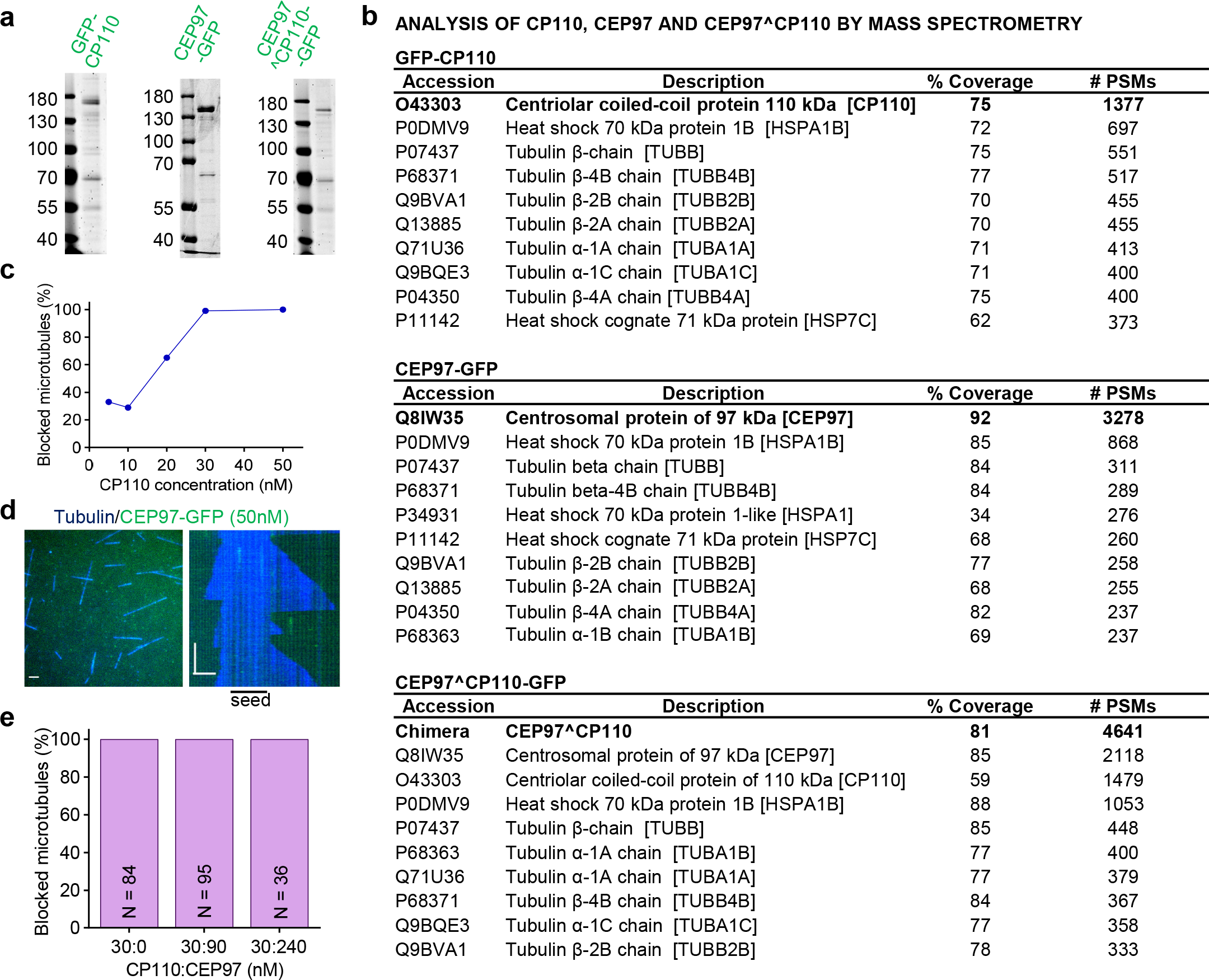
Characterization of purified GFP-CP110 and CEP97. (a) SDS-PAGE of GFP-CP110, CEP97-GFP and CEP97^CP110-GFP, purified from HEK293T cells. Gels were stained with Coomassie Brilliant Blue. (b) Analysis of purified GFP-CP110, CEP97-GFP and CEP97^CP110-GFP by mass spectrometry. (c) The proportion of fully blocked MTs with increasing concentrations of GFP-CP110 in in vitro reconstitution assays. The number of analyzed MTs was 91, 28, 142, 105 and 140 MTs at 5, 10, 20, 30 and 50 nM, respectively. (d) A still image and kymograph representing dynamic MT behavior in the presence of CEP97-GFP. Bars, 2 µm (horizontal); 1 min (vertical). (e) Bars plot showing CEP97-GFP does not affect the plus-end blocking of dynamic MTs in vitro by GFP-CP110. The numbers of MTs evaluated are indicated on the bar plots.

**Supplementary Figure S2.**
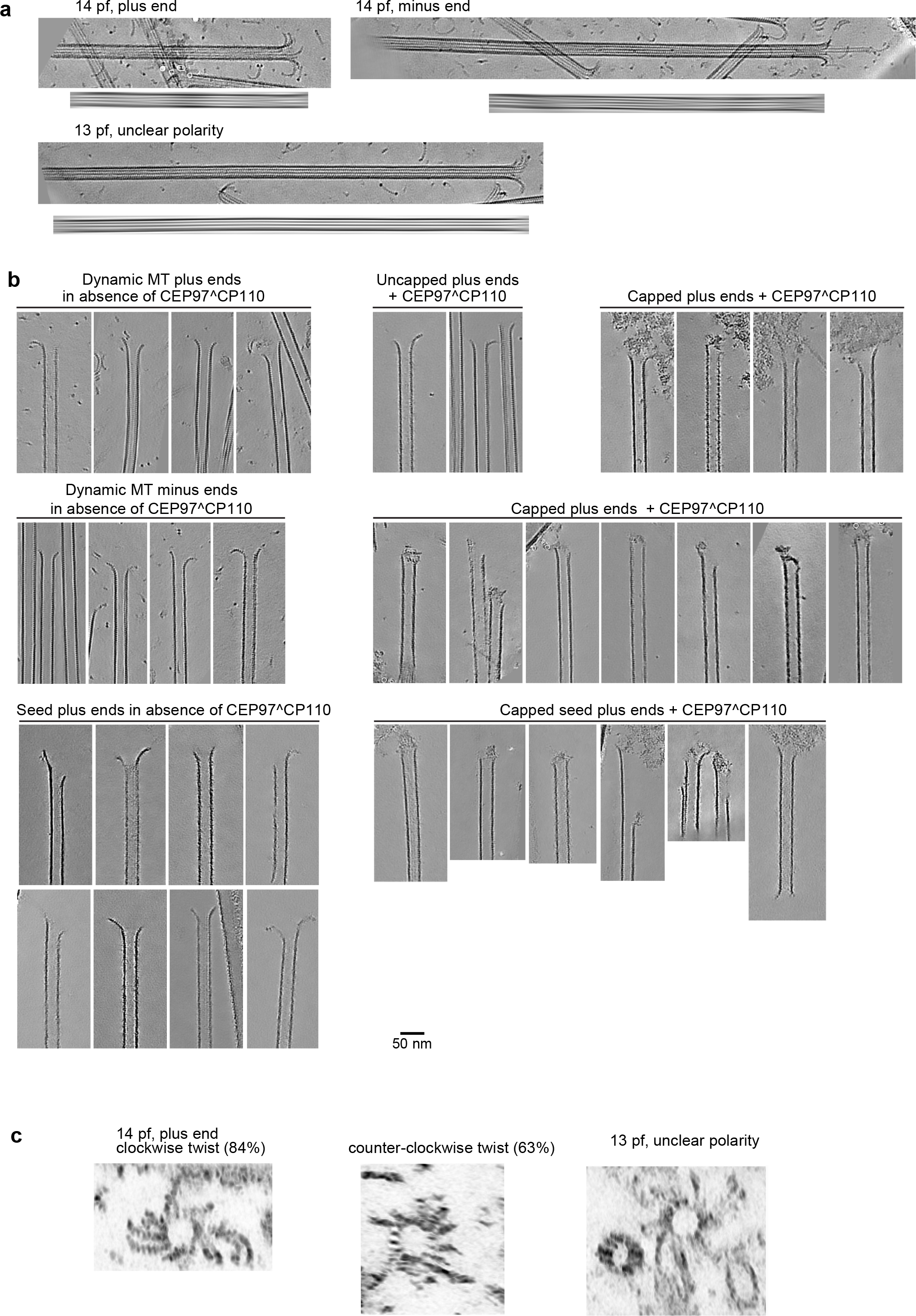
Characterization of MT ends by cryoET. (a) Determination of MT polarity. For each MT: sum of slices containing the MT (top), and the same image Fourier-filtered at origin (bottom). (b) Gallery of MT ends. Scale bar: 50 nm. (c) Sum of slices obtained from the tomograms rotated 90 degrees to illustrate the end-on view of protofilament flares. Plus ends typically show clockwise twist pattern, while minus ends typically show counter-clockwise patter. The twist pattern is also observed for 13-protofilament MT ends.

**Supplementary Figure S3.**
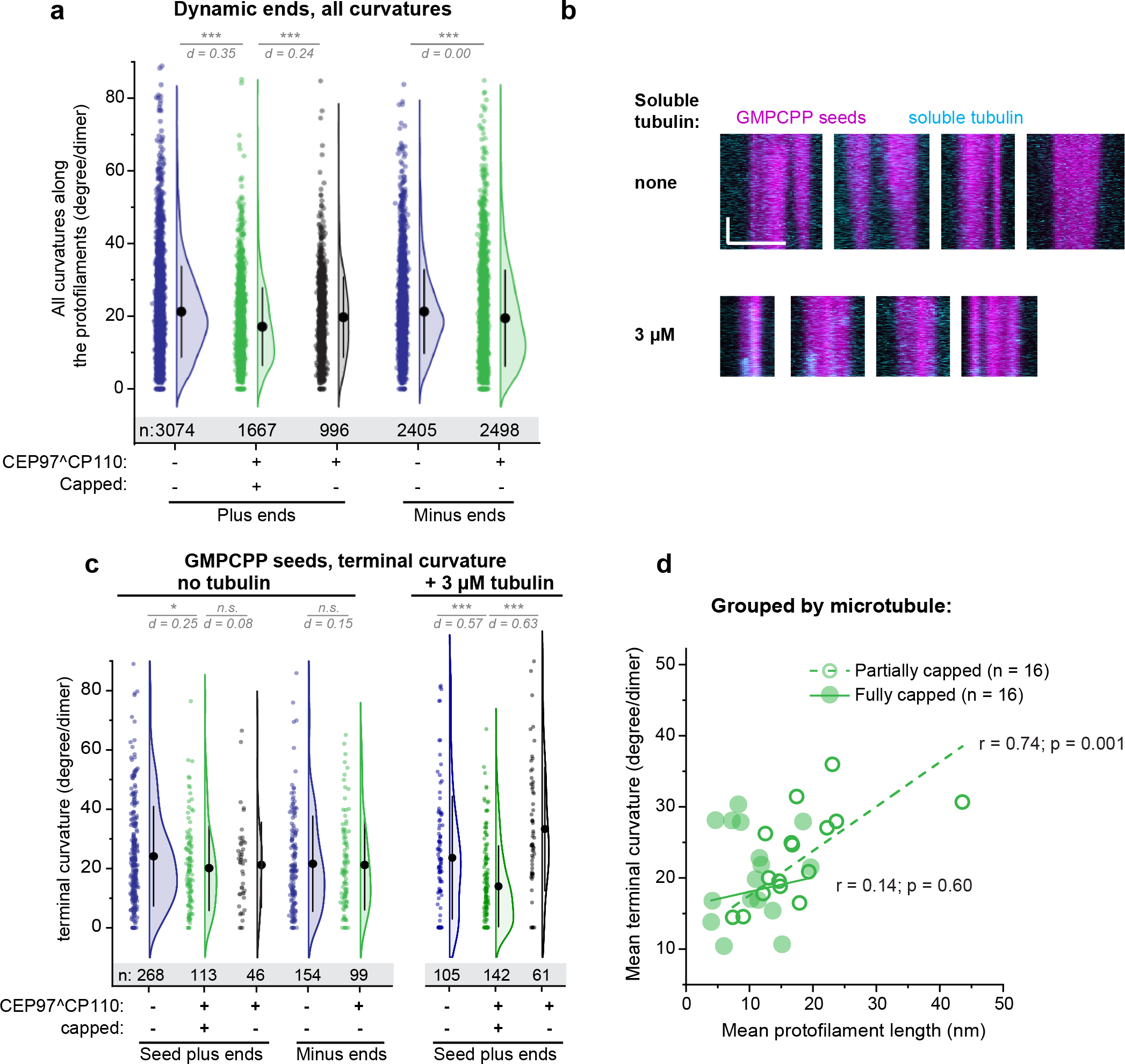
Characterization of MT ends by cryoET. (a) Distribution of all curvatures along protofilaments with non-zero length, obtained in the presence of soluble tubulin. Statistical summary: d indicates effect size (Cohen’s d) expressed in units of SD; ***, P< 0.001 (Mann-Whitney test). (b) Kymographs showing GMPCPP-stabilized seeds polymerized using HiLyte488 tubulin (magenta) in absence (top row) or presence (bottom row) of soluble tubulin labelled with TMR (cyan). Scale bars: vertical (60 s), horizontal (5 µm). (c) Distribution of all protofilament curvatures obtained from samples of GMPCPP seeds without soluble tubulin or with 3 µM soluble tubulin. Statistical summary: d indicates effect size (Cohen’s d) expressed in units of SD; ns – no significant difference; *, P<0.05, ***, P< 0.001(Mann-Whitney test). (d) Correlation between average terminal curvature and average protofilament length per MT plus end. *r* – Pearson correlation coefficient, *p* – probability that the slope of the correlation is different from zero.

**Supplementary Figure S4.**
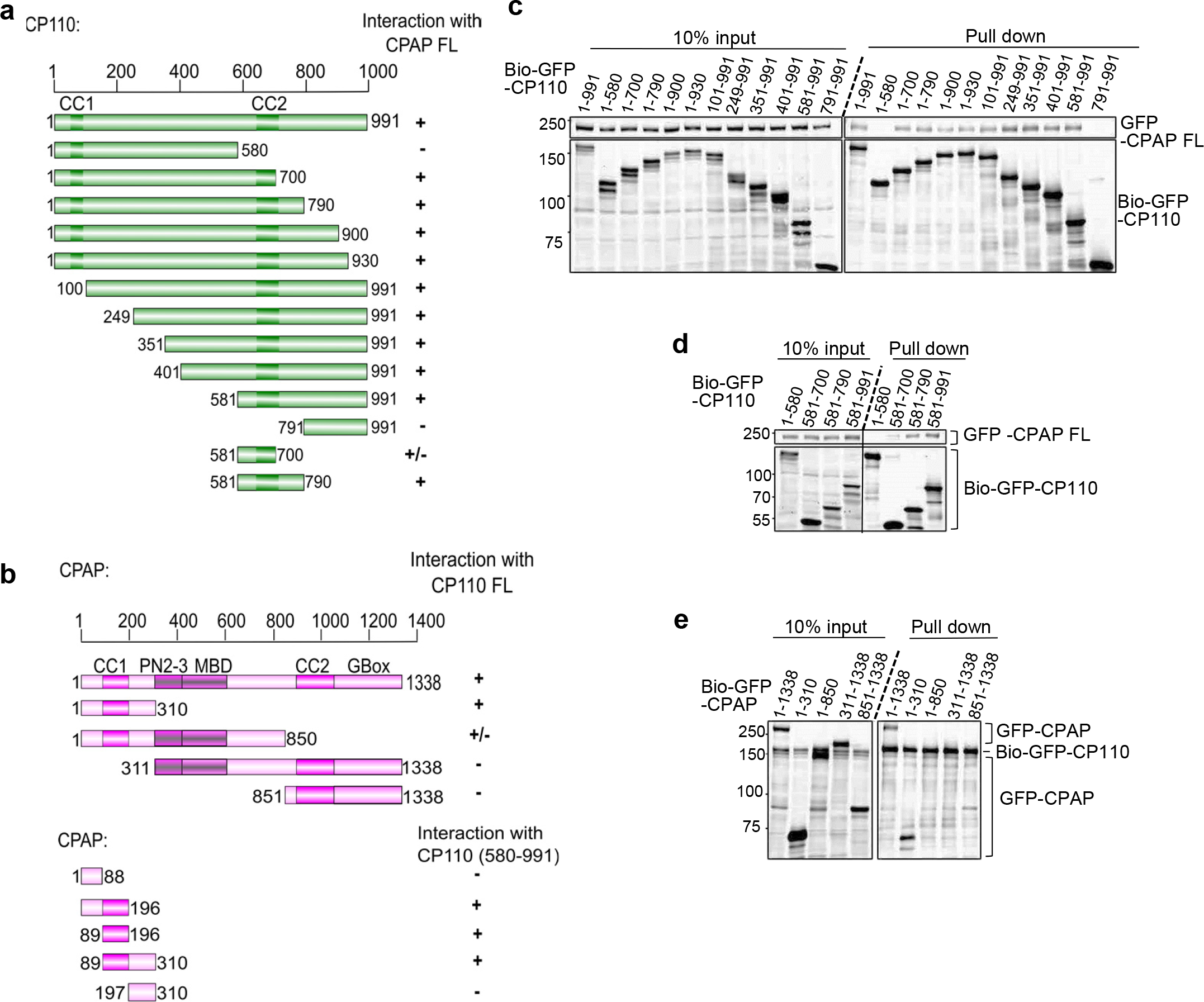
Mapping of the interaction between CP110 and CPAP. (a, b) Schemes of CP110 and CPAP illustrating the deletion mutants used in this study. CC, coiled coil; PN2-3, tubulin-binding domain; MBD, MT-binding domain; “+”, interaction between CPAP and CP110; “-”, no interaction between CPAP and CP110. (c, d) Streptavidin pull-down assays with BioGFP-CP110 truncations as bait and full-length GFP-CPAP as prey. (e) Streptavidin pull-down assays with BioGFP-CPAP truncations as bait and full-length GFP-CP110 as prey. All the assays were performed with extracts of HEK293T cells co-expressing the indicated constructs and BirA and analyzed by Western blotting with anti-GFP antibodies.

**Supplementary Figure S5.**
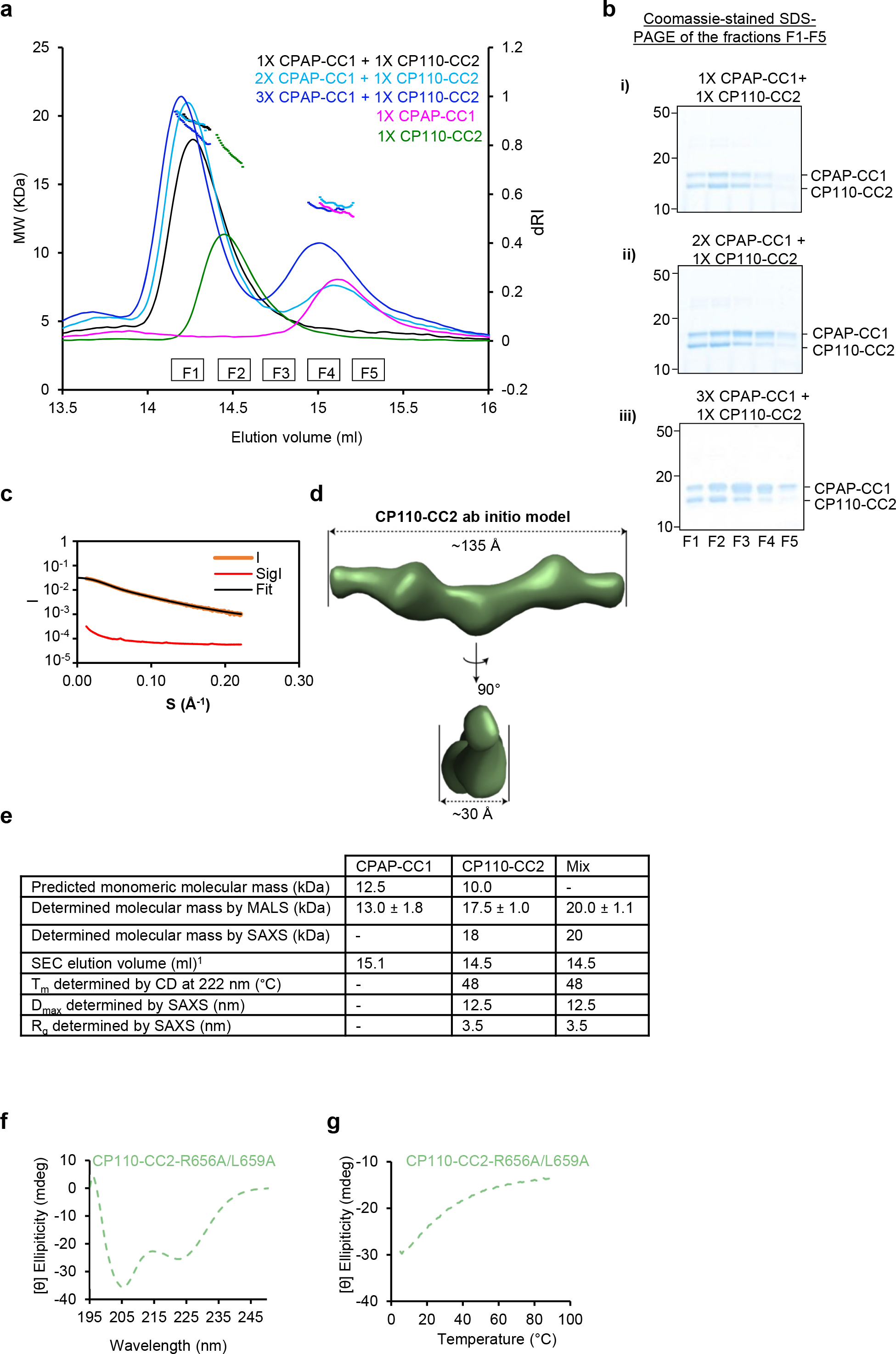
Biophysical characterization of CPAP-CC1, CP110-CC2, and CPAP-CC1-CP110-CC2. (a) SEC-MALS analyses of CPAP-CC1 (magenta lines) and CP110-CC2 (green lines) alone, and mixtures of CPAP-CC1 with CP110-CC2 at molar ratios of 1:1 (black line), 2:1 (light blue line), and 3:1 (dark blue line). (b) Coomassie-stained SDS-PAGE of the fractions F1-F5 indicated in panel (a) and collected from SEC-MALS runs obtained with mixtures of CPAP-CC1 and CP110-CC2. SDS-PAGE analysis of the elution peak fractions centered at around 14.3 ml (corresponding to the molecular weight of CPAP-CC1/CP110-CC2 heterodimer) of the various mixtures revealed equally intense protein bands corresponding to CPAP-CC1 and CP110-CC2. These results support the idea that two CPAP-CC1 monomers react with one CP110-CC2 dimer to form two stable heterodimers in solution. (c, d) SAXS analysis of the CP110-CC2 homodimer. (c) Solution X-ray scattering intensity over scattering angle from CP110-CC2. The fit to the data yielding the interatomic distance distribution is shown with a black line. (d) Surface representation of the X-ray scattering volume of CP110-CC2, at 30 ± 2 Å estimated precision, derived from averaging 22 particle models calculated by *ab initio* fit to the scattering data. (e) Table summarizing biophysical parameters of CPAP-CC1, CP110-CC2 and an equimolar mixture of CPAP-CC1 and CP110-CC2 obtained by SEC-MALS, CD, and SAXS. (f, g) CD spectrum (f) recorded at 15°C and thermal unfolding profiles (g) recorded by CD at 222 nm of CP110-CC2 R656A/L659A (light green dashed lines).

**Supplementary Figure S6.**
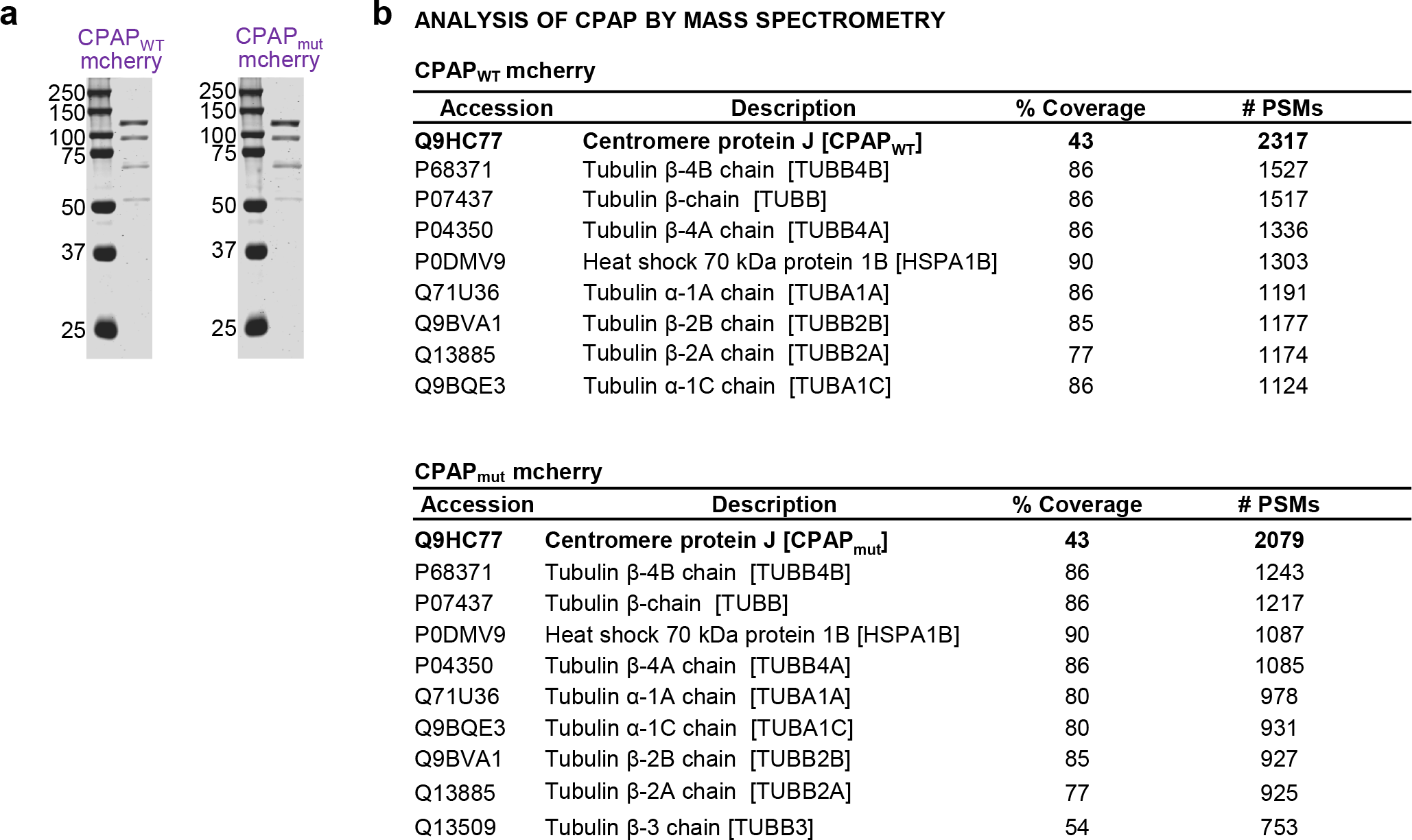
Characterization of purified CPAP proteins. (a) SDS-PAGE of CPAPWT and CPAPmut, purified from HEK293T cells. Gels were stained with Coomassie Brilliant Blue. (b) Analysis of purified GFP-CP110, CEP97-GFP and CEP97^CP110- GFP by mass spectrometry.

**Supplementary Figure S7.**
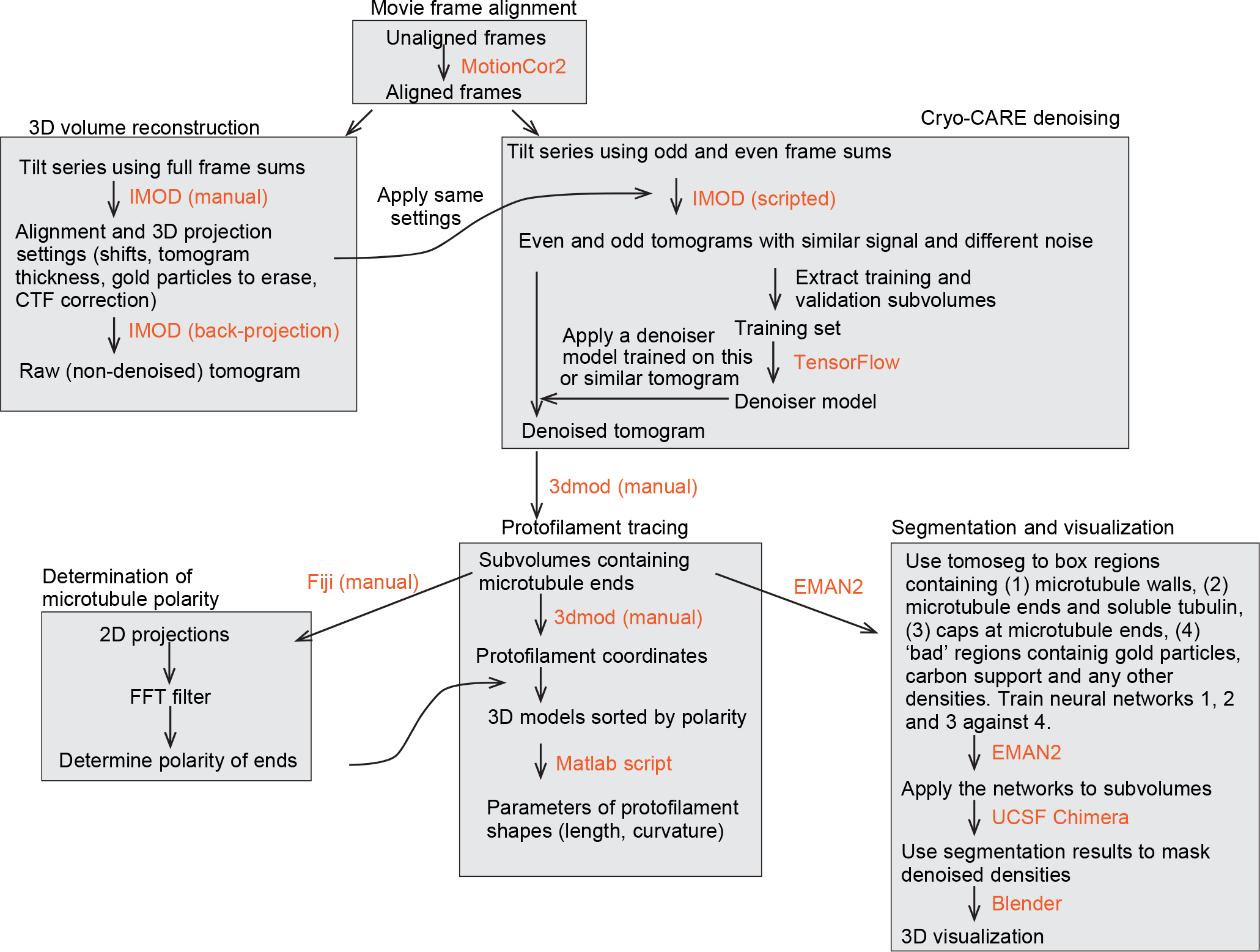
Schematic flow-chart illustrating the pipeline for 3D reconstruction, denoising, segmentation and visualization of tomographic volumes.

Supplementary Video S1. 3D view of MT plus ends in the absence and presence of CEP97^CP110.

The video shows MT plus ends in the absence (left) or presence (right) of CEP97^CP110-GFP. The denoised densities were segmented into tubulin and MTs (blue) and all other densities (green) as described in Methods. Manually segmented models with coordinates of tubulin protofilaments for each of the plus ends are shown in orange.

